# Novel mechanistic insights into the role of Mer2 as the keystone of meiotic DNA break formation

**DOI:** 10.1101/2020.07.30.228908

**Authors:** Dorota Rousova, Vaishnavi Nivsarkar, Veronika Altmannova, Saskia K. Funk, Vivek B. Raina, David Liedtke, Petra Janning, Franziska Müller, Heidi Reichle, Gerben Vader, John R. Weir

## Abstract

In meiosis, DNA double strand break (DSB) formation by Spo11 initiates recombination and enables chromosome segregation. Numerous factors are required for Spo11 activity, and couple the DSB machinery to the development of a meiosis-specific ?axis-tethered loop? chromosome organization. Through in vitro reconstitution and budding yeast genetics we here provide architectural insight into the DSB machinery by focussing on a foundational DSB factor, Mer2. We characterise the interaction of Mer2 with the histone reader Spp1, and show that Mer2 directly associates to nucleosomes, likely highlighting a contribution of Mer2 to tethering DSB factors to chromatin. We reveal the biochemical basis of Mer2 association with Hop1, a HORMA domain-containing chromosomal axis factor. Finally, we identify a conserved region within Mer2 crucial for DSB activity, and show that this region of Mer2 establishes an interaction with the DSB factor Mre11. In combination with previous work, we establish Mer2 as a keystone of the DSB machinery by bridging key protein complexes involved in the initiation of meiotic recombination.

## Introduction

Meiotic recombination is one of the defining features of eukaryotic sexual reproduction. In addition to creating the genetic diversity that fuels speciation and evolution, meiotic recombination fulfils a direct mechanistic role in establishing connections between initially unpaired homologous chromosomes. Meiotic recombination is initiated by programmed double-strand DNA break (DSB) formation by the transesterase Spo11^1^. Meiotic DSBs are preferentially repaired via recombination from the homologous chromosome which, depending on how recombination intermediates are processed, can yield crossovers (reviewed in^2^). Together with sister chromatid cohesion, crossovers provide the physical linkage between homologous chromosomes which is necessary to ensure meiotic faithful chromosome segregation. In most organisms, crossover formation is associated with, and influenced by synapsis between homologs, established by the assembly of the synaptonemal complex. The formation of meiotic DSBs by Spo11 needs to be carefully orchestrated and controlled. In addition to Spo11, at least ten additional factors are required for Spo11-dependent DSB activity, and collectively these factors are referred to as the meiotic DSB machinery. Functional and biochemical analysis has begun to reveal the logic of the assembly of the DSB machinery. A picture is emerging in which several distinct subcomplexes are co-recruited into Spo11-activity proficient chromosomal foci. In addition to the core DSB machinery, several other factors promote meiotic DSB activity. For example, DSB formation occurs in the context of a distinctive chromatin loop-axis architecture which is established concomitantly with the entry of cells into the meiotic program (Figure 1A). In budding yeast, this proteinaceous axis is made up of a meiosis-specific cohesin complex (Rec8-cohesin) in combination with the coiled-coil scaffolding protein Red1, and the HORMA domain protein Hop1^3^. In cells lacking meiotic axis components, Spo11-dependent DSB formation is severely impaired (but not completely abolished, as in DSB machinery mutants), and efficient recruitment of meiotic DSB factors depends on axis establishment. In addition, DSB placement and formation is influenced by histone modifications (specifically Histone H3-K4 methylation, which in budding yeast cells directs Spo11 to gene promotor regions). These nucleosomal interactions of the DSB machinery are proposed to occur with genomic regions which are located in the chromatin loop that emanate away from the chromosome axis (to which the DSB machinery is tethered).

**Figure 1.**
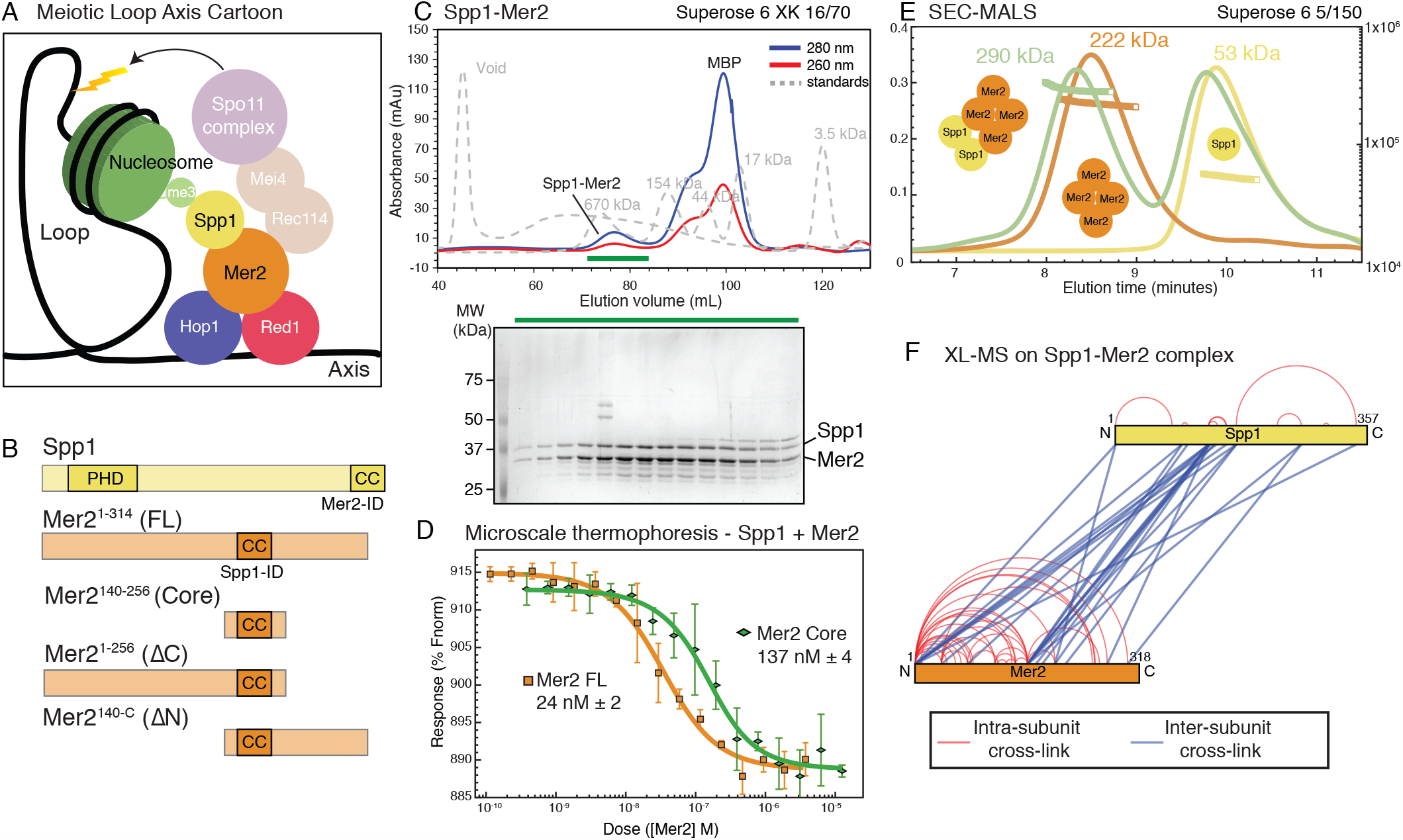
Spp1 binds to Mer2 tetramerisation domain with a 2:4 stoichiometry A) Cartoon of meiotic loop-axis architecture and the role of Mer2. During meiosis proteins Red1 and Hop1 form a protein:DNA (coloured black) axis together with cohesin (not shown for clarity). Loops of chromatin are extruded from the axis, where DNA breaks are made by the Spo11 complex (pale magenta). Breaks are directed to the proximity of H3K4me3 nucleosomes (green) through the combined activity of Mer2 and Spp1. Mer2 (orange) is also thought to interact with additional Spo11 accessory proteins (Rec114 and Mei4, pale orange). B) Domain diagram of Mer2 and Spp1. The four principal Mer2 constructs used throughout this study are shown. The only clear feature of Mer2 (and its orthologs IHO1 in mammals, Rec15 in fission yeast and Prd3 in plants) is the central coiled-coil motif. Spp1 is shown for comparison with its N-terminal PHD domain and C-terminal Mer2 interaction domain which is predicted to contain a coiled-coil. C) Purification of Mer2-Spp1 complex. A complex of Mer2 and Spp1 was purified to homogeneity. The MBP tags on Spp1 and on Mer2 were cleaved prior to loading. Degradation products of Mer2 full-length protein are also visible. Molecular weight markers are shown in grey. The relative absorbance of the complex at 280 nm and 260 nm shows that it is free of any significant nucleic acid contamination. The selected fractions (green line) were loaded onto an SDS-PAGE gel and stained with InstantBlue. D) Microscale thermophoresis of Mer2-Spp1. Two different Mer2 constructs (Mer2^FL^, orange squares; Mer2^core^ green diamonds) were titrated against Red-NHS labelled untagged Spp1 (20 nM constant concentration), and the change in thermophoresis was measured. Experiments were carried out in triplicate and the *Kd* was determined from the fitting curve. E) SEC-MALS of different Spp1-Mer2 samples. Measured molecular masses are indicated Three illustrative SEC-MALS experiments are shown for Spp1 (yellow)[theoretical mass of monomer = 42 kDa], MBP-Mer2^core^ (orange) [theoretical mass of a tetramer = 226 kDa] and a complex of Spp1 bound to MBP-Mer2^core^ (green) [theoretical mass of 2:4 complex = 310 kDa].. F) XL-MS of Mer2-Spp1 complex. Full-length Spp1 and Mer2 were used, and in both cases the overhang remaining after cleaving the N-terminal fusion proteins was present. Samples were cross-linked with DSBU, and data analysis carried out according to the materials and methods. Cross-links were filtered so as to give a 1% false discovery rate. Figure was prepared using XVis ^55^. Intramolecular cross-links shown in red, intermolecular cross-links in blue.

A key component of the meiotic DSB machinery is Mer2. This protein (also known as Rec107) was originally identified as a high-copy number suppressor of the *mer1* phenotype (Mer1 was later shown to be a co-factor for the splicing of various meiotic mRNAs, including Mer2^4^), after which it was shown to be essential for meiosis ^5^. Mer2 is central to the temporal control of Spo11-dependent DSB formation, being the target of S-Cdk and DDK (Cdc7-Dbf4) phosphorylation that presumably allows the binding of the Spo11-associated factors Rec114 and Mei4 to Mer2^6–8^. This regulation plays a crucial role in the spatiotemporal assembly of the DSB machinery. In addition to forming a complex with Rec114-Mei4, Mer2 interacts directly with the PHD domain-containing protein Spp1 ^9,10^. Spp1 binds to nucleosomes that are tri- (or di-) methylated on H3K4 (*i*.*e*. H3K4^me3^ nucleosomes)^11,12^, and this association is important for the association of Spo11-dependent DSB formation with gene promoter regions. Spp1 is canonically part of the COMPASS (a.k.a. Set1 complex), but during meiosis, Spp1 forms an independent, and mutually exclusive, interaction with Mer2. The reciprocal interaction domains between Spp1 and Mer2 have been previously identified ^9,10^. The C-terminal region of Spp1 interacts with a central, predicted coiled-coil, region of Mer2^9,10^(Figure 1B). A single amino acid substitution in Mer2 (V195D) is sufficient to disrupt the interaction with Spp1 (as judged by Yeast-2-Hybrid (Y2H) analysis)^13^. Through its interaction with Spp1, a key role for Mer2 appears to link the Spo11-machinery directly to Spp1-mediated nucleosome interactions. Interestingly, Spp1 associated with Mer2 has a longer residence time on nucleosomes when compared with Spp1 when part of COMPASS^14^, suggesting additional functions for Mer2 in mediating nucleosome tethering. In line with the central position for Mer2 in DSB machinery assembly is the observation that - in contrast to deletion of Set1 or Spp1 which severely reduces, but not eliminates DSB formation - *mer2*Δ cells completely fail to form meiotic DSBs^15^. Mer2 likely establishes additional biochemical interactions that establish a functional Spo11-assembly. For example, homologs of Mer2 in fission yeast^16^ and mouse^17^ interact with meiotic chromosome axis-associated HORMA proteins, suggesting that Mer2 can establish a link between the chromosome axis (via HORMA protein interaction), and chromatin loops (through Spp1 association).

Despite hints to the central position of Mer2 in assembly of DSB machinery, a more comprehensive biochemical understanding of these interactions is critically needed. Here we use a combination of *in vitro* biochemical reconstitution with yeast genetics to investigate several distinct protein-protein interactions of Mer2. We examine the interaction of Mer2 with Spp1, nucleosomes, with proteins of the meiotic axis, and with additional members of the DSB machinery. Our results report a more complete picture of Mer2 as a foundational component of the meiotic DSB machinery, including novel functions, and provide mechanistic explanations for a number of previously observed phenomena revolving around the regulation of meiotic DSB formation.

## Results

### Mer2-Spp1 is a constitutive complex with a 2:4 stoichiometry

We first focussed on the described interaction between Mer2 and Spp1. In order to probe the various possible functions of Mer2, we made use of four principal expression constructs, the full-length protein (residues 1-314 from hereon abbreviated to “Mer2^FL^”), Mer2 amino acids 1-256, lacking the C-terminal 58 residues (from hereon “Mer2^C^”), Mer2 residues 140 to 314 (from hereon “Mer2^N^”) and Mer2 containing residues 140-256 (from hereon referred to as “Mer2^core^”) (Figure 1B). Previous work has identified the Spp1 interaction region to be contained within Mer2^core^ (specifically Mer2 residues 165-232^10^). Using our in-house expression system, “InteBac”^18^, we produced full-length Spp1 and all Mer2 proteins in *E*.*coli* with N-terminal MBP tags to facilitate protein solubility. We could successfully remove the MBP-tag using the 3C protease from both Spp1 and Mer2, though in the case of Mer2^N^ and Mer2^Core^ the 3C cleavable MBP tag could not be removed, presumably due to steric hindrance by the MBP tag that precludes efficient cleavage. In co-lysis experiments, we found that we could purify a complex of Mer2 and Spp1 to homogeneity, and free of nucleic acid contamination (Mer2^FL^ with Spp1 shown as an example in Figure 1C, note the apparent A^260^ to A^280^ ratio as evidence of a lack of nucleic acid contamination), thus indicating that the interaction between Mer2 and Spp1 does not require any PTMs or additional cofactors and that the interaction was robust enough to survive extensive co-purification in >300 mM NaCl.

Using microscale thermophoresis (MST) we measured the binding affinity of Spp1 to Mer2^FL^ (Figure 1D orange trace, squares) and Spp1 to Mer2^Core^ (Figure 1D green trace, diamonds). Spp1 bound Mer2^FL^ with a *K*_*D*_ of 24 nM (+/-2), and to Mer2^Core^ with a *K*_*D*_ of 137 nM (+/-4). Mer2 constructs lacking the “core” showed comparatively weak binding (Supplementary Figure 1). Thus, we confirm that the majority of the Spp1 binding interface is indeed within the core of Mer2, as reported earlier^10,13^, but there does appear to be some contribution to Spp1 binding provided by the N- and C-terminal regions of Mer2. We measured the molecular mass of Mer2 by size exclusion chromatography coupled to multi-angle light scattering (SEC-MALS) and concluded that Mer2^core^ is the tetramerisation region (Figure 1E, orange trace and Supplementary Figure 2A-D), consistent with recent observations of Mer2{^19^. Interestingly, while Mer2^Ccore^ is monomeric (Supplementary Figure 2D), Mer2^Ncore^ is dimeric (Supplementary Figure 2C) indicating the presence of a dimerisation region in the C-terminal region of Mer2 between residues 255 and 314, which presumably aids in the stability of a full antiparallel coiled-coil tetramer, since we observe no species larger than a tetramer.

Given that the tetramerisation region of Mer2 is also the principal Spp1 binding region, we determined the stoichiometry of the Mer2-Spp1 complex. First, we determined that full-length Spp1 alone is a monomer (Figure 1E yellow trace, Supplementary Figure 2 E and F). We next analysed the stoichiometry of Mer2:Spp1 complexes. We measured the size of a complex of MBP-Mer2^core^ with Spp1 (Figure 1E green trace) and determined its mass to be 290 kDa. The theoretical mass of a 2:4 (Spp1:Mer2) complex is 310 kDa, whereas a 1:4 (Spp1:Mer2) complex is 268 kDa. Given the possible ambiguity in determination of stoichiometry we also measured complexes of MBP-Mer2^FL^ together with Spp1 (Supplementary Figure 2G) which gave complex sizes best fitting a 2:4 (Spp1:Mer2) stoichiometry. Taken together we conclude that the Mer2 tetramer binds two copies of Spp1, establishing a complex in a 4:2 stoichiometry. Thus we show a novel function for Mer2 in not simply binding Spp1, but importantly, in mediating the dimerisation of Spp1. In light of the inherent twofold symmetry of nucleosomes, we suggest that this 4:2 constellation might aid in the recognition of modified nucleosome tails by Spp1.

Next we probed the structural organisation of Spp1-Mer2 further using cross-linking coupled to mass spectrometry (XL-MS) using the 11Å-spacer crosslinker disuccinimidyl dibutyric urea (DSBU) (Figure 1F). XL-MS of Spp1-Mer2 revealed that, while the “core” of Mer2 showed many cross-links with Spp1, these were, unexpectedly, not with the previously described C-terminal interaction domain “Mer2-ID” of Spp1^10^. Instead the Mer2 core showed numerous cross-links with a region of Spp1 immediately C-terminal to the PHD domain. Furthermore there were additional cross-links between the N- and C-terminal regions of Mer2 and Spp1, consistent with the residual binding affinity we observed in MST (Supplementary Figure 1) The intramolecular cross-linked pattern of Mer2 alone (Supplementary Figure 3A) was very similar in the presence and absence of Spp1. As such we can likely exclude a significant structural rearrangement of Mer2 upon association with Spp1. We also compared the cross-linking pattern observed previously for Mer2 alone^20^ (Supplementary Figure 3B). This revealed that the pattern was broadly similar with a mixture of long- and short-distance cross-links, best explained by an antiparallel arrangement of the four Mer2 polypeptides within the coiled-coil core of the complex, as recently suggested^19^. One striking difference is the extensive cross-links emanating from the N-term of our Mer2. The most likely explanation is that the overhang remaining on our Mer2 preparation after removal of the N-terminal fusion protein is four amino acids longer, and thus more flexible.

### Spp1-Mer2 complex binding to H3K4me3 modified mononucleosomes

In order to study the role of the Spp1-Mer2 complex binding to H3K4^me3^ nucleosomes, we created synthetic H3K4^me3^ mononucleosomes. Briefly, we mutated K4 of Histone H3 to cysteine whereas the single naturally occurring cysteine of the natural H3 sequence was mutated to alanine (C110A). H3C4 was converted to H3K4^me3^ using a trimethylysine analogue as previously described{^21^. We then reconstituted H3K4^me3^ into octamers, and subsequently into mononucleosomes using 167bp Widom sequences (see Materials and Methods for further details). Due to the dimeric nature of nucleosomes, and because our reconstitutions showed a 2:4 Spp1-Mer2 complex stoichiometry, we hypothesised that dimerization of Spp1 would presumably lead to more tight binding to H3K4^me3^ nucleosomes, and set out to test this idea. We compared the apparent binding affinity of Spp1, GST-tagged Spp1 (which mediated dimerisation) and the Spp1-Mer2 complex. We observed that the Spp1:Mer2 assembly (in which two copies of Spp1 are present) bound more tightly to nucleosomes, as compared to monomeric Spp1 alone (which bound relatively weakly to nucleosomes, consistent with the reported 1 µM affinity of the PHD dopared the observed binding of Spp1:Mer2 with GST-Spp1, we found that GST-Spp1 exhibited an intermediate apparent binding affinity. A potential corollary of this observation is that Mer2, in addition to triggering Spp1 ‘dimerization’ might directly contribute to nucleosome binding.

We established that the Spp1-Mer2 complex was capable of forming a stable complex with H3K4^me3^ nucleosomes in solution (Figure 2C). Note that we do not observe a complete shift of nucleosomes; we suspect that this is due to not having an optimal buffer condition (in this experiment we have tried to balance the buffer conditions required for Mer2 (high salt), Spp1 (Zn^2+^ ions for the PHD domain) and nucleosomes (EDTA)). As our complex is currently not suitable for high-resolution structural studies, we made use again of XL-MS to determine a topological architecture of the Spp1-Mer2-H3K4^me3^ mononucleosome complex (Figure 2D). We detected many more internal Spp1 cross-links than observed in the Spp1-Mer2 complex alone (Figure 1F), suggesting that either the binding to nucleosomes brings the two Spp1 moieties (in the 2:4 complex) closer to one another, or that there is an internal rearrangement of domains of Spp1. Most strikingly however, the cross-linking revealed that most of the cross-links between the Spp1-Mer2 complex and the nucleosomes are via regions of Mer2, in the N-term, core and C-term regions. This observation strength-ened the idea that Mer2 might directly contribute to nucleosome binding. We modelled the location of the Spp1-Mer2 cross-links onto the previously determined structure of a mononucleosome^22^ (Figure 2E). We observe that the cross-links cluster around histone H3, but more generally around the DNA entry/exit site on the nucleosomes. These observations suggest a large Mer2:Nucleosome interface, and a surprisingly smaller Spp1:Nucleosome interface.

**Figure 2.**
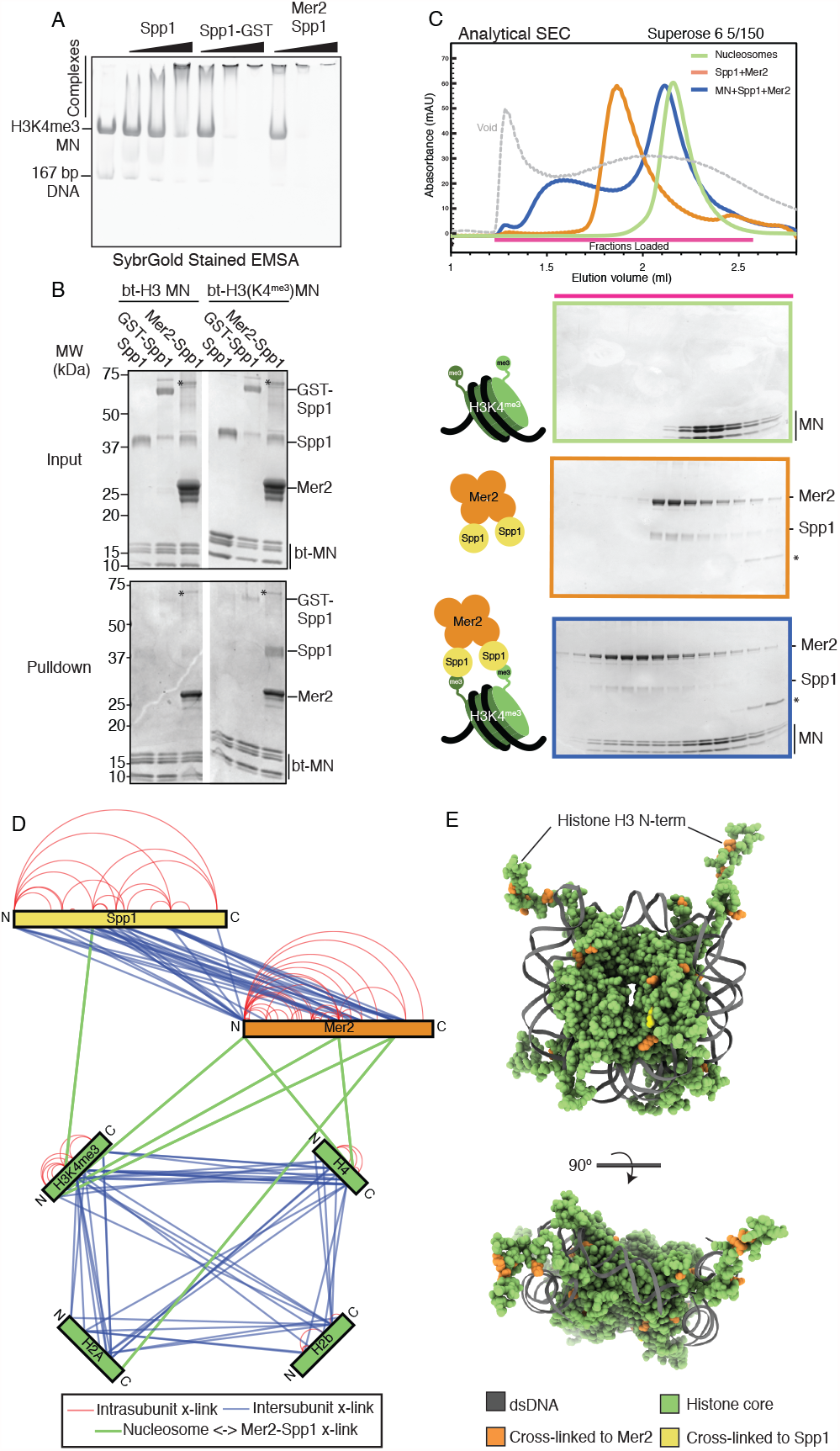
Spp1-Mer2 complex binding to H3K4^me3^ mononucleosomes A) EMSA of different Spp1 variants on H3K4me3 nucleosomes. 0.2 µM of H3K4me3 nuleosomes were incubated with 0.33, 1 and 3 µM protein (Spp1, Spp1 with a C-terminal GST fusion and Mer2-Spp1). Gel was post-stained with SybrGold. B) Biotinylated-Mononucleosome pulldown of different Spp1 variants. 0.5 µM nucleosomes wrapped with 167 bp of biotinylated DNA (either with or without the H3K4me3 modification) were incubated with 1.5 µM protein (same proteins as in a)). Samples were taken for the input before incubation with streptavidin beads. The beads were then washed and eluted with 1x Laemmli buffer. Input and elution samples were run on a 10-20% SDS-PAGE gel and stained with InstantBlue. Asterisk marks residual uncleaved MBP-Mer2 in the Mer2-Spp1 lanes. C) SEC analysis of Spp1-Mer2-MN complex. 50µL of 5 µM H3K4me3 mononucleosomes (green), 50 µL of 5µM Mer2-Spp1 complex (orange) and 50 µL of a 1:1 (5µM of each) mixture (blue) were run on a Superose 6 5/150 column. The same fractions were loaded in each case (magenta line) onto an SDS-PAGE gel and stained with InstantBlue. D) XL-MS analysis of the Spp1-Mer2-Mononucleosome complex. Samples were cross-linked with DSBU, and data analysis carried out according to the materials and methods. Cross-links were filtered so as to give a 1% false discovery rate. Figure was prepared using XVis <u>^55^</u>. Intramolecular cross-links are shown in red, intermolecular cross-links shown in blue. Cross-links between Mer2 and nucleosomes, and Spp1 and nucleosomes shown as thick green lines. E) Model of nucleosome cross-links. The nucleosomes proximal cross-links from D) were modelled onto a crystal structure of a *X. laevis* nucleosome (PDB ID 1KX5^22^. Those histone residues that cross-linked to Mer2 are coloured in orange, those that cross-link to Spp1 in yellow. A side and top down view of the nucleosome are provided. DNA is coloured in dark grey, and histone residues that did not cross-link to Spp1 or Mer2 are coloured green.

### Mer2 binds directly to nucleosomes with a 4:1 stoichiometry

Our pulldown data, SEC experiments and XL-MS data all suggested that Mer2 might bind to mononucleosomes directly, perhaps providing additional affinity to the Spp1:nucleosome interaction. If true, this would be a previously unreported function of Mer2. We tested whether Mer2^FL^ could bind to *unmodified* mononucleosomes in a SEC experiment, and found that it surprisingly formed a stable complex (Figure 3A). Given that Mer2 is a tetramer that binds two copies of Spp1, and given that Mer2 has been previously shown to form large assemblies on DNA^19^ we asked what the stoichiometry of a Mer2-Mononucleosome complex was. To do this, we first used Mass Photometry (MP), a technique that determines molecular mass in solution at low concentrations based on the intensity of scattered light on a solid surface^23^. In MP, we observe a mix of three species, free Mer2 tetramer (measured at 127 kDa), free mononucleosomes (measured at 187 kDa) and a complex at 303 kDa (Figure 3B). The experiment was carried out at a protein concentration of 60 nM, which suggests that the dissociation constant *(K*_*D*_) is somewhat less than 60 nM (at *K*_*D*_ under equilibrium one would observe 50% complex formation, we observe less than 50% in Figure 3B). We next asked whether Mer2 might form larger assemblies on nucleosomes as higher concentrations. Using SEC-MALS (Figure 3C), we observed a complex of 341.0 kDa which nearly perfectly matches a theoretical complex consisting of one Mer2 tetramer plus one mononucleosome (340 kDa, summarised in Figure 3D). Additionally we observe a small fraction of a very high molecular weight assembly (though not an aggregate) of 10.96 MDa. This could be an oligomer of Mer2, of Mer2 on nucleosomes or Mer2 on free DNA. It has recently been reported that Mer2 binds directly to DNA^19^. Given that we observe Mer2 cross-links at the nucleosome DNA entry/exit site we therefore tested whether Mer2 might simply be binding the free DNA ends on mononucleosomes. Using analytical EMSAs we found that Mer2 binds with a 6-fold higher apparent affinity to nucleosomes (5 nM vs. 30 nM), than to the same 167 bp DNA used to reconstitute the nucleosomes (Figure 3E). The discrepancy between *K*_*D*_ determined by EMSA and the apparent *K*_*D*_ from MP is presumably because EMSAs are non-equilibrium experiments, carried out by necessity at very low salt^24^.

**Figure 3.**
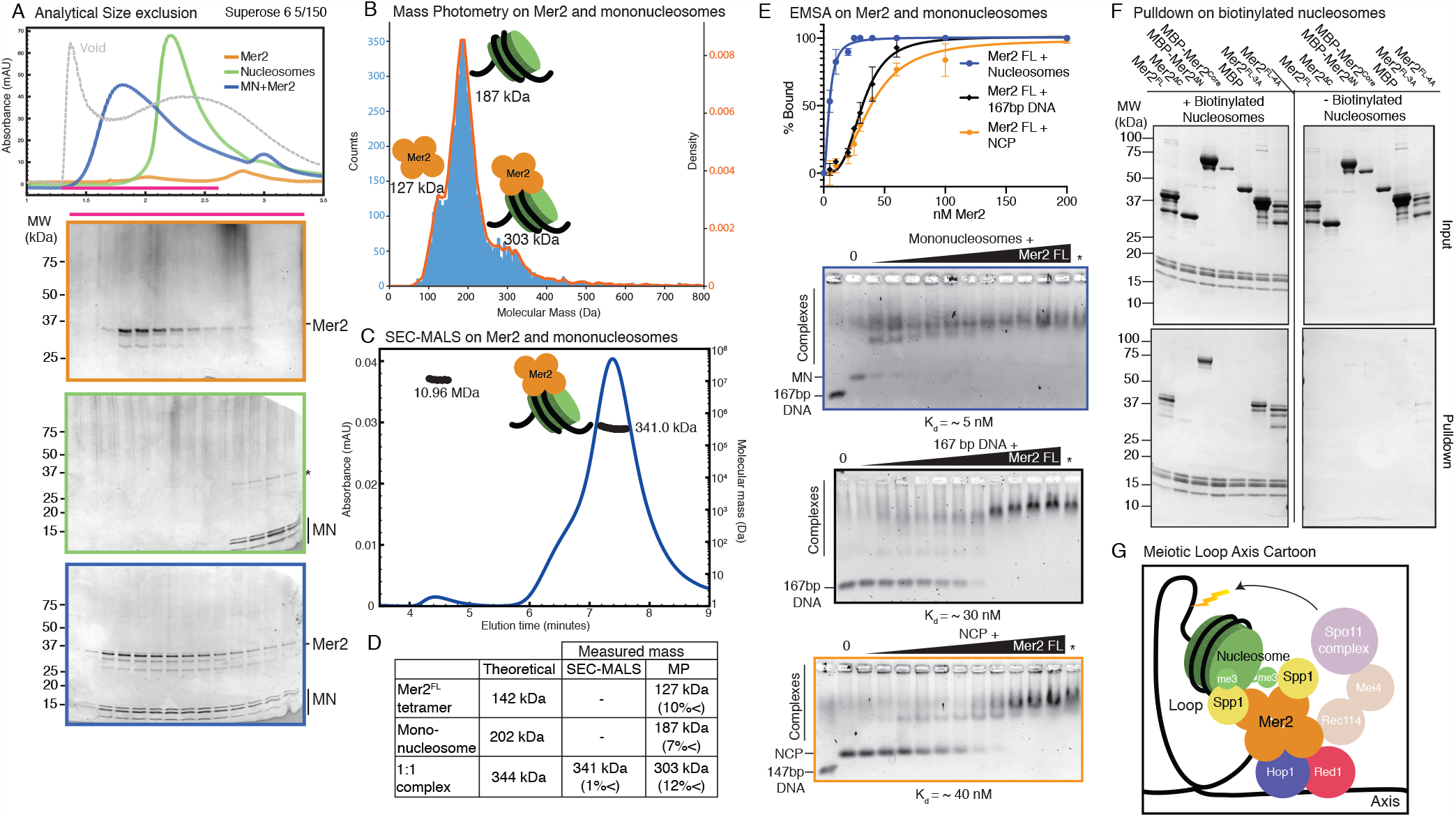
Mer2 binds directly to nucleosomes. A) SEC analysis of Mer2-MN complex. 50 µL sample of 5 µM mononucleosomes (green), 50 µL of 5 µM full length untagged Mer2 (orange) and a mixture of Mer2 and mononucleosomes (blue) were run on a Superose 6 5/150 column. The same fractions were loaded in each case (magenta line) onto an SDS-PAGE gel and stained with InstantBlue. B) Mass Photometry of Mer2 and mononucleosomes. 60 nM of Mer2 and mononucleosomes were mixed and analyzed using a Refeyn One mass photometer. Three separate species were identified and the molecular mass determined using a molecular mass standard curve created under identical buffer conditions. Negative data points (i.e. unbinding events) were excluded. C) SEC-MALS on Mer2-MN complex. Full-length, untagged Mer2 (Mer2^FL^) was incubated with mononucleosomes and subject to size exclusion chromatography coupled to multi angle light scattering (SEC-MALS). Absorbance at 280 nm was constantly monitored (blue trace) Two distinct species were observed, one at 341 kDa, and another at 10.96 MDa (black trace). D) Summary of molecular mass values for Mer2FL, mononucleosomes (with 167 bp DNA) and a 1:1 complex. E) EMSAs of untagged Mer2 FL. Mer2 was titrated against a constant 5 nM concentration of mononucleosomes (blue), 167 bp “601” DNA (black), or nucleosome core particle (orange). Binding curves were derived based on the Mer2 dependent depletion of free nucleosomes, DNA or NCP, and based on four independent experiments with error bars indicating the SD. Asterisk denotes the background SYBR-Gold staining from the highest protein concentration alone. F) Pulldowns of Mer2 constructs with nucleosomes. Biotinylated nucleosomes (left panel) were incubated with different Mer2 constructs (as indicated), and samples taken for the input gel. The complexes were captured using Streptavadin beads, washed, and eluted in 1x Laemmelli buffer for the pulldown gel. A control experiment (right panel) was conducted without biotinylated nucleosomes to measure non-specific Mer2 interaction with the streptavidin beads. G) Cartoon summarising our findings and changes to the model so far. Mer2 binds to two Spp1 subunits (yellow), which in turn bind to a double H3K4^me3^ modified nucleosome. Mer2 (orange) makes direct contacts with the nucleosome (green), presumably via the nucleosomal DNA.

Next we asked what effect using a smaller length of DNA to reconstitute nucleosomes might have. We reconstituted histone octamers on 147 bp DNA (from now on referred to as nucleosome core particles (NCP)^25,26^). We found that with no free DNA ends Mer2 did bind with a lower affinity to NCPs, but nonetheless still with an apparent *K*_*D*_ of ∼40 nM (Supplementary Figure 4A). We then asked whether mutating the common binding site, the “acidic patch” on H2A (E56T-E61T-E64T-D90S-E91T-E92T)^27^ might have an effect on Mer2 binding. In order to enhance potential differences in binding, this was also done on NCPs. Mer2 bound to NCP acidic patch mutants essentially as well as wild-type NCPs (Supplementary Figure 4A). We then asked whether Mer2 might be recognising the histone tails. We therefore prepared “tailless” NCPs (see materials and methods). Surprisingly Mer2 bound very tightly to tailless NCPs with an apparent K_d_ of ∼5 nM (Supplementary Figure 4A). We suggest that this might indicate that under the conditions of an EMSA, the histone tails are shielding either the histone cores or the NCP DNA and interfering with binding by Mer2.

We next asked which region of Mer2 might be involved in binding nucleosomes (reconstituted with 167 bp DNA). Initially we carried out EMSAs with Mer2^core^, Mer2^N^ and Mer2^C^. Mer2^C^ and Mer2^N^ appeared to bind slightly weaker than Mer2^FL^ (K_d_ ∼12.5 nM and ∼30 nM respectively), whereas Mer2^core^ showed no binding at all (Supplementary Figure 4B). Apparently there was equal contribution to nucleosome binding from both termini. In order to refine this further, we carried streptavidin pulldown using mononucleosomes reconstituted with biotinylated DNA against different Mer2 constructs. This approach had the advantage of being able to use a more physiological, and as such more stringent, buffer. We confirmed that the Mer2^core^ did not bind nucleosomes, but neither did Mer2^C^, strongly suggesting that the main nucleosome interaction region of Mer2 lies in the C-terminal 58 amino acids (Figure 3F). We propose that Mer2, in addition to enabling the ‘dimerization’ of Spp1 via its central tetramerization domain, provides a direct binding interface with nucleosomes (Figure 4G), with this functionality possibly being encoded in its C-terminal region.

**Figure 4.**
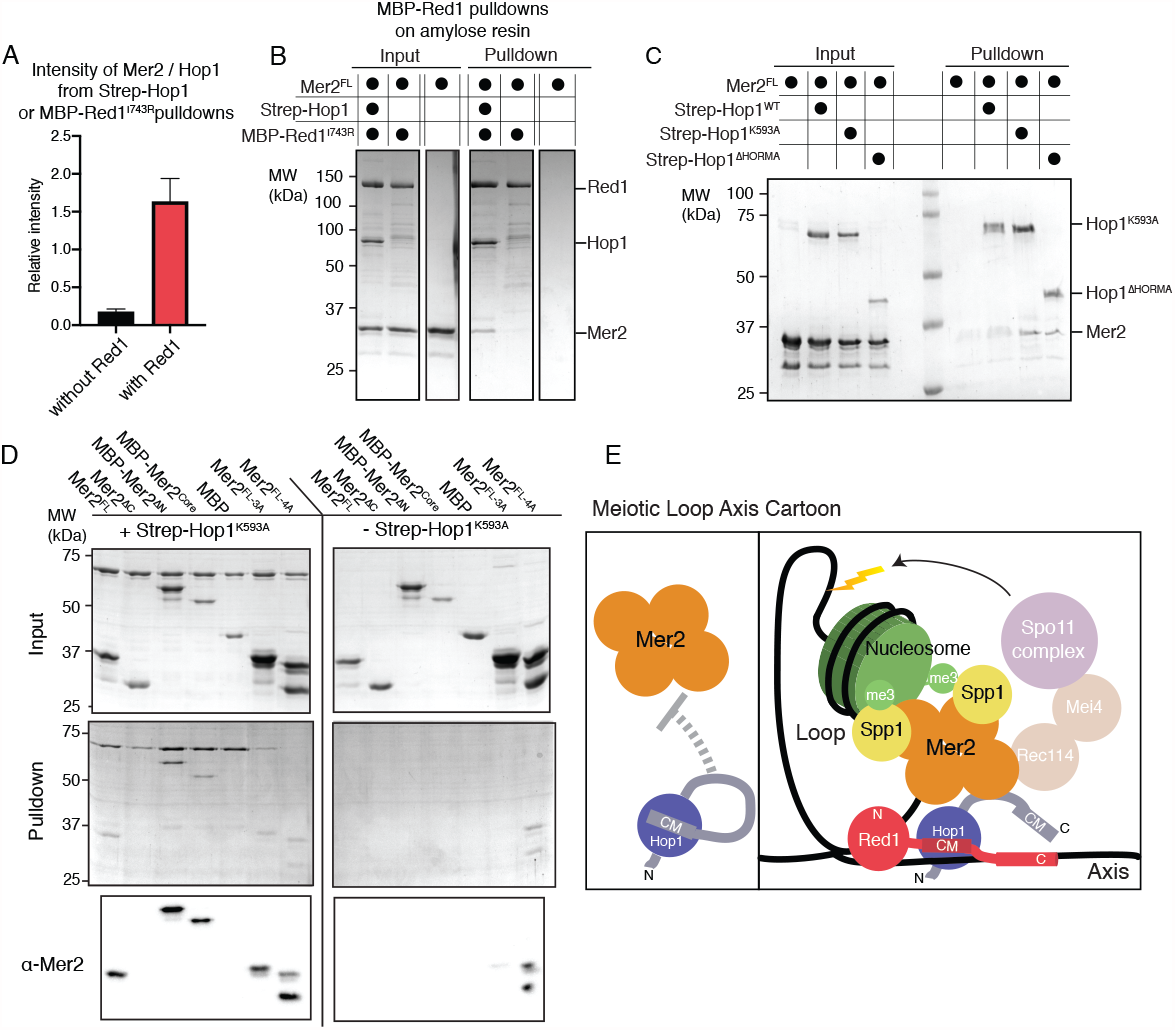
Mer2 binds to the axis via open Hop1 A) Quantification of Streptactin pulldowns on either Strep-Hop1 (black) or the Strep-Hop1/Red1^I743R^-MBP (red) complex against Mer2 (from gels shown in Supplementary Figure 5). In all cases the intensity of Mer2 was quantified as a factor of Hop1 intensity. Error bars are based on the SD from three independent experiments. B) Amylose pulldown of the co-expressed Red1^I743R^-MBP/Hop1 complex and Red1^I743R^-MBP alone against Mer2. Pulldown was carried out as described in materials and methods. In the rightmost lane Mer2 alone was used as a negative control on the beads. C) Streptactin pulldown of different 2xStrepII-Hop1 constructs (as indicated) against untagged Mer2^FL^. N-terminally 2xStrep-II tagged Hop1 constructs were purified as indicated and used as a bait against Mer2 full-length. Pulldown was carried out as described in the materials and methods. D) Streptactin pulldown of Strep-Hop1^K593A^ as bait to capture different Mer2 constructs as indicated. The rightmost pulldown is the control showing the non-specific binding of different Mer2 constructs to the Streptactin beads. An anti-Mer2 western was carried out (lower panel) to confirm the specific interactions due to the weakness of the Mer2 bands in a coomassie stained gel. E) Cartoon representation of the meiotic loop axis structure. In the left panel Hop1 that is bound to its own CM cannot bind to Mer2. In the right panel Hop1 that is bound to the CM of axial Red1 releases its own CM and is then able to bind to Mer2.

### Mer2 binding of Hop1 requires “unlocking” the C-terminus of Hop1

In addition to Spp1, several lines of evidence point to an association between Mer2 and meiotic HORMA domain-containing factors. In fission yeast and mouse, the functional homologs of Mer2 (named Rec15 and IHO1, respectively), have been shown to interact with Hop1/HORMAD1^16,17^. Likewise in budding yeast Mer2 exhibits a very similar chromatin association pattern to Hop1^28^. Meiotic HORMA proteins are integral members of the meiotic chromosome axis, and are needed to recruit Mer2 to chromosomes^28,17^. Hop1 (like most other known HORMA domains) can exist in two topological states (‘open/unbuckled’ (O/U) or ‘closed’ (C)), in which the closed state can embrace a binding partner via a closed HORMA-closure motif ‘safety belt’ binding architecture^29^ The closure motif (also referred to as CM) is a loosely conserved peptide sequence encoded in HORMA binding partners. The meiotic HORMA proteins are unique among HORMA proteins in the fact that they contain a CM at the end of their own C-terminus (Hop1 residues 585-605^29^ endowing these factors with the ability to form an intramolecular (closed) HORMA-CM configuration. The association of Hop1 with chromosomes is mediated by an interaction with Red1 which depends upon a similar CM:HORMA-based interaction. The CM of Red1 (located 340-362^29^ binds to Hop1 with a higher affinity than the closure motif of Hop1^29^ There is mounting evidence in budding yeast and *Arabidopsis*, that, in addition to a chromosomal pool, a significant pool of Hop1 is non-chromosomal (in the nucleoplasm or cytoplasm)^30,31^. Once bound to its own CM, Hop1 will not be able to interact with its chromosomal axis binding partner Red1. Due to the high local concentration of the intramolecular CM, free Hop1 is expected to rapidly transition from the ‘open/unbucked’ into the intramolecular ‘closed’ state^29^. The closed state (whether it is intramolecular or, for example, with the CM of Red1) can be reversed by the action of the AAA+ ATPase Pch2/TRIP13^32,33^. As such, within the nucleoplasm/cytoplasm, Pch2/TRIP13 activity serves to generate enough ‘open/unbuckled’ Hop1 that is proficient for incorporation into the chromosomal axis, via a CM-based interaction with Red1. On the other hand, when recruited to chromosomes, that same Pch2/TRIP13 activity is expected to dismantle Hop1-Red1 assemblies, as such leading to removal of Hop1 from chromosomes^34,35^.

We tested the ability of Mer2 to interact with the proteins of the meiotic axis Hop1 and Red1, with a focus on Hop1. Initially, we purified Hop1 using an N-terminal Twin-Strep-II tag and used it to pulldown Mer2. We observed a faint band corresponding to Mer2 in the Hop1 pulldown, indicative of a weak interaction (Supplementary Figure 5, lane 1). Note that based on the high relative concentration of the CM in Hop1, this Hop1 is expected to largely consist of (intramolecular) closed Hop1. We then co-expressed Red1-MBP containing a I743R (from hereon referred to as Red1^I743R^-MBP) mutation with Hop1 in insect cells. The I743R mutation should prevent Red1 from forming filaments, but still allow Red1 to form tetramers^36^. Given that we have an excess of Hop1 in our Hop1-Red1 purification, we carried out a pulldown on the MBP-tag of Red1 (using amylose beads) with Mer2 as prey. In this case, we observed considerably more Mer2 binding, when measured relative to Hop1, its putative direct binding partner (Supplementary Figure 5, lane 3). We quantitated the Mer2 intensity relative to the Hop1 band in both pulldowns, and from three independent experiments, and observed a ∼6-fold increase in Mer2 binding (Figure 4A). We reasoned that this difference could either be due to Red1 interacting directly with Mer2, or that Red1 induces a conformational change in Hop1 that facilitates Mer2 binding. To test the former idea we purified Red1^I743R^-MBP in the presence or absence of Strep-Hop1. Using amylose affinity beads to capture the MBP moiety of Red1 we tested the capture of Mer2 both in the presence and absence of Hop1 (Figure 4B). We could detect no Mer2 interaction when pulling on Red1^I743R^-MBP in the absence of Hop1. As such, we conclude that Mer2 does not have significant affinity for Red1. These data argue that Hop1, when bound to Red1, is acting as an efficient recruiter of Mer2 to this complex. How could this increased affinity of Hop1 for Mer2 be influenced by association with Red1? Based on the known biochemical basis of the Hop1-Red1 interaction, we can imagine that the interaction of Red1 with Hop1 “releases” the C-terminus of Hop1 which could create a “chain” of Hop1 moieties (akin to what has been observed in *C. elegans*^37^) or could simply liberate the C-terminus of Hop1 for binding to non-self partners. We propose that such configurations would create or expose Mer2-specific binding interfaces that are shielded when Hop1 is bound intramolecularly to its own C-terminal CM. The second model makes two testable predictions: 1) the region of Hop1 that interacts with Mer2 should be encoded in its (non-HORMA) C-terminus, and 2) impairing intramolecular CM-HORMA binding should ‘unlock’ the binding ability of Hop1 with Mer2, regardless of Red1 presence. To test these predictions, we created two additional Hop1 constructs, one where the N-terminal HORMA domain is missing (Hop1^HORMA^) and another where the conserved lysine in the closure motif has been mutated to alanine (Hop1^K593A^) disrupting the ability of the CM of Hop1 to interact with its HORMA domain thus forcing Hop1^K593A^ into an “unlocked” state^29^. Both Hop1^HORMA^ and Hop1^K593A^ were purified with an N-terminal 2xStrep-II tag (as for Hop1^WT^) and their ability to bind to Mer2 was tested (Figure 4C). In line with the idea that binding of Hop1 to Red1 could enable Mer2 association via a direct competition with intramolecular CM-HORMA binding, we observed robust Mer2 binding for both Hop1^256-C^ and Hop1^K593A^ but not Hop1^WT^.

Having determined that an “unlocked” Hop1 is necessary for Mer2 interaction, we then asked which region of Mer2 is necessary for Hop1 interaction. Using N-terminally 2xStrepII tagged Hop1^K593A^ as bait, we queried a variety of Mer2 constructs using Streptactin beads (Figure 4D). Due to the weak staining of Mer2 under these conditions we also carried out an anti-Mer2 western blot (Figure 4D lower panel). We determined that Hop1 was capable of interacting, apparently equally well, with all Mer2 constructs, except for Mer2^C^. Why would the Mer2^core^ be capable of binding to Hop1, but the C not? The answer may lie in the complex antiparallel arrangement of the Mer2 coiled-coils and thus N- and C-termini of Mer2 relative to one another and to the core of Mer2. As such one could imagine that the interaction region for Hop1 lies in the core of Mer2, which might be shielded by the N-terminus when the C-terminus of Mer2 is not present. These results hint at a complex regulation, on the Mer2 side, underlying Mer2-Hop1 interaction, and we believe that high-resolution structural studies of the Mer2-Hop1 interface should eventually be able to provide deeper insight into this interaction. Taken together we propose a new model for Mer2 recruitment to the meiotic axis in which a “locked” Hop1 cannot bind to Mer2 (Figure 4E, left) but once incorporated into Red1, Hop1 is “unlocked” and recruits Mer2 to the meiotic axis via an interaction region present in its exposed C-terminus (Figure 4E, right).

### Conserved N-terminal motif in Mer2 is essential for spore formation

Due to the potential difficulties in assigning defined interaction regions within Mer2 when using truncation constructs, we aimed to obtain separation of function alleles by introducing selected point mutants instead. Making use of sequence alignments from (evolutionary closely and more distantly related) Mer2 orthologs, we identified a previously undescribed conserved region, in the N-terminal domain between amino acids 52 and 71 (Figure 5A, Supplementary Figure 6). This particular stretch of amino acids stands out in the protein sequence of Mer2 (and homologs) since, in addition to the central coiled-coil region, this region is one of the few regions which shows sequence similarities across evolutionary distant species (such as yeast and human). To probe a potential function of this conserved region, we created two different alleles, which we here refer to as *mer2-3a* and *mer2-4a*. In *mer2-3a*, we mutated 3 conserved residues *W58, K61*, and *L64* to alanine (Figure 5A). *mer2-4a* contains the following mutations that fall within the same domain: *D52A, E68A, R70A* and *E71A*. We integrated plasmids carrying C-terminally 3HA-tagged versions of wildtype *MER2, mer2-3a*, or *mer2-4a* (note that all these constructs are driven by *pMER2*) in *mer2* strains. All three constructs lead to comparable expression levels of Mer2 during meiotic prophase, although we note different mobility of the Mer2^3A^-3HA or Mer2^4A^-3HA as compared to Mer2-3HA, which might indicate altered post-translational modifications, such as phosphorylation (Figure 5B). Alternatively, it might reflect an inherent effect of the introduced mutations (compare for example also the migrating patterns of wild type Mer2 with Mer2^3A^ on SDS-PAGE from our *in vitro* preparations, where meiosis-specific post-translational modifications are presumably absent (see Figure 6B).

**Figure 5.**
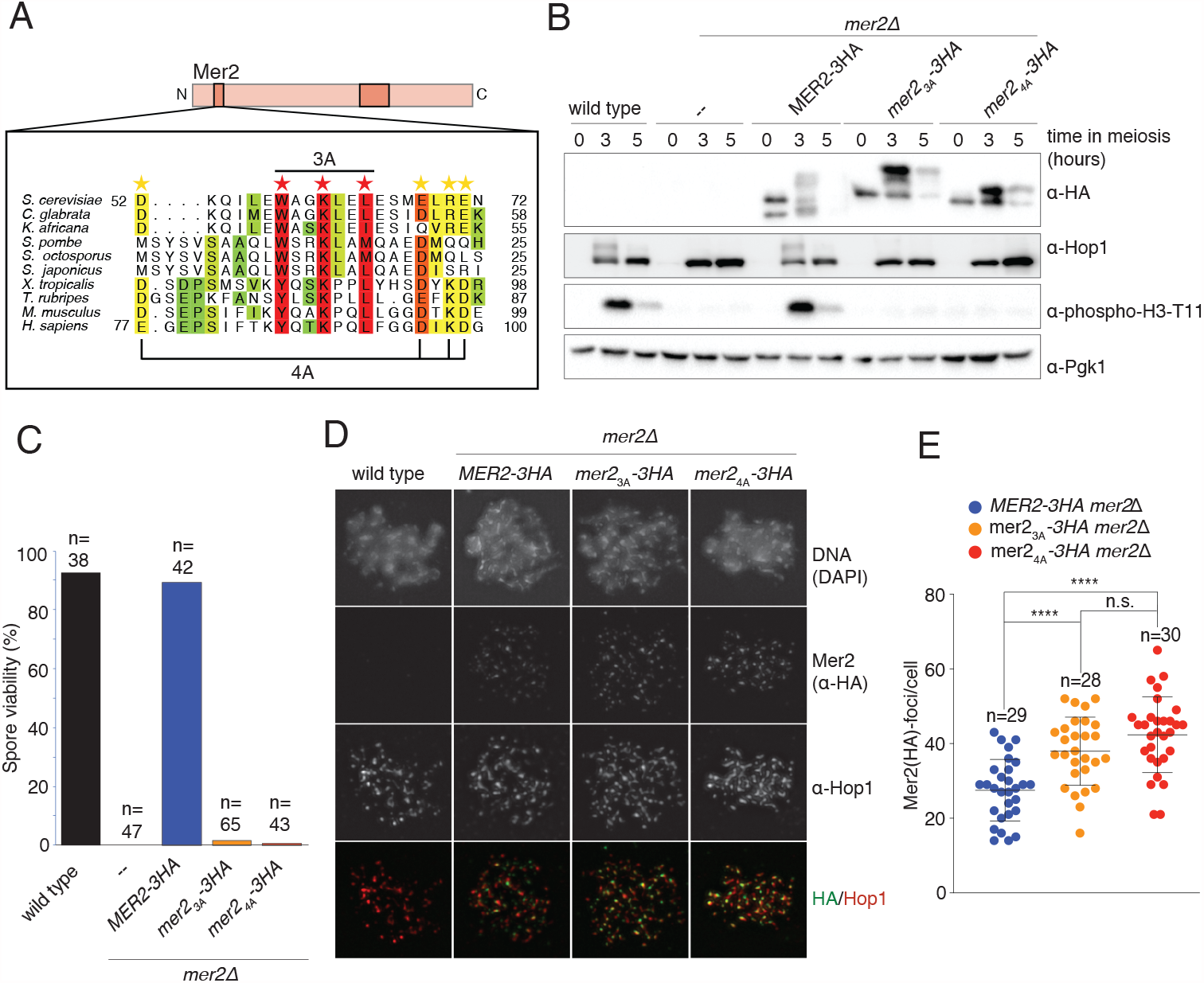
Mer2 3A and 4A mutations prevent DSB formation A) Sequence alignments of Mer2 orthologs from multiple clades revealed a previously undescribed conservation in the N-terminal region. We created two mutants, the “3A” mutant (W58A, K61A, L64A; red stars) based on three universally conserved residues and the “4A” mutant (D52A, E68A, R70A, E71A; yellow stars). B) Western blot analysis of meiotic yeast cultures of wildtype, *mer2*D, *MER2-3HA mer2*D, *mer2-3a-3HA mer2*D, and *mer2-4a-3HA mer2*D *strains*. Time of induction into the meiotic program indicated. See Supplementary Table for strain information. Pgk1 was used as a loading control. C) Quantification of spore viabilities of wildtype, *mer2*D, *MER2-3HA mer2*D, *mer2-3a-3HA mer2*D, and *mer2-4a-3HA mer2*D strains. Number of dissected tetrads is indicated. See Supplementary Table for strain information. C) Representative images of meiotic chromosome spreads stained for Mer2 (*–*-HA (green), Hop1, and DNA (blue) using DAPI from wildtype, *MER2-3HA mer2*D, *mer2-3a-3HA mer2*D, and *mer2-4a-3HA mer2*D strains at t=3 hours after induction into the meiotic time course. D) Quantification of the number of Mer2 foci on chromosome spreads of strains used in (D). Mean and standard deviation are indicated. p 0.05, p 0.01, and p 0.0001; n.s. (non-significant) > 0.05. Mann-Whitney U test. Number of analyzed cells is indicated.

**Figure 6.**
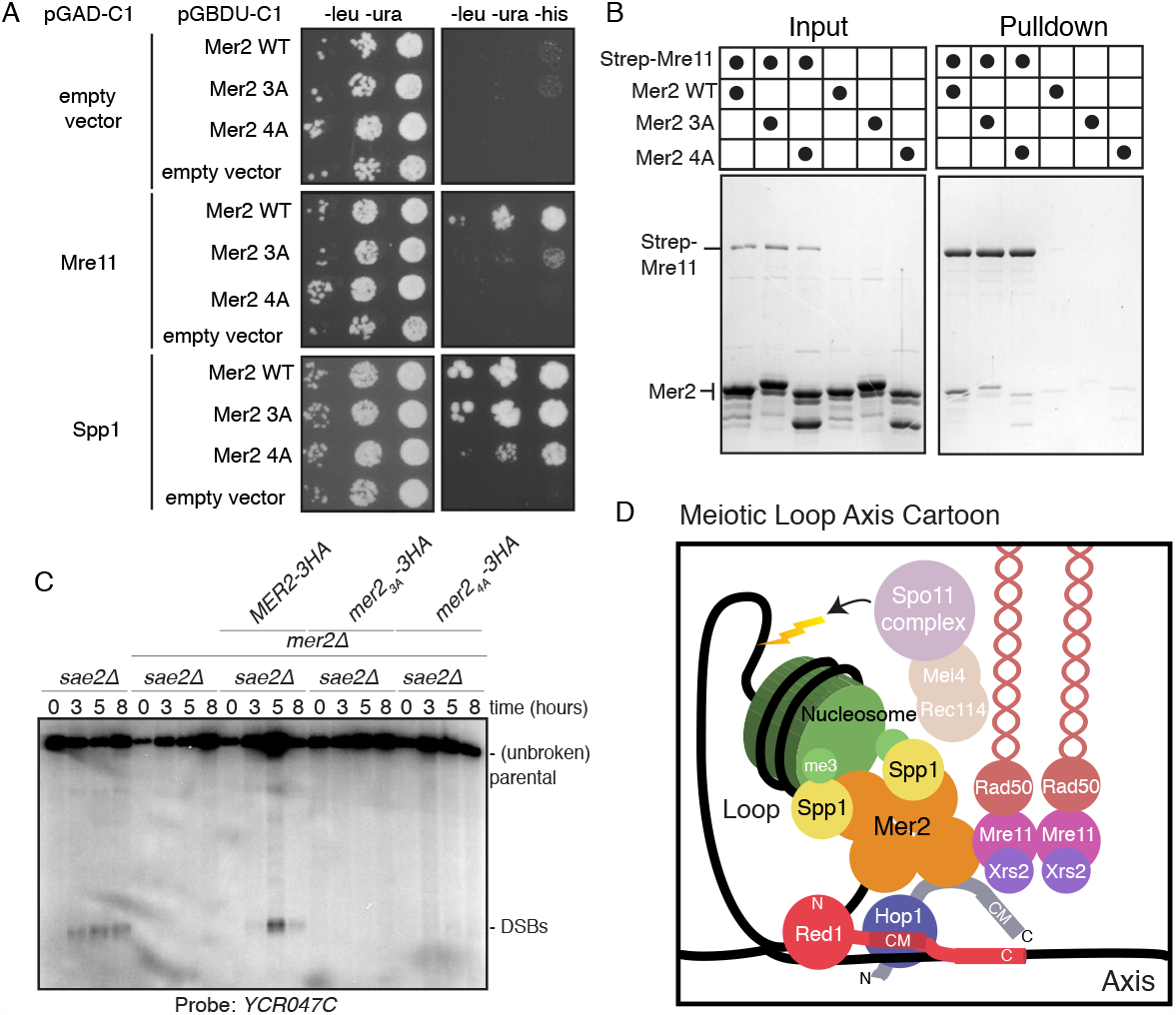
Mer2 binds directly to Mre11 A) Yeast-two-hybrid experiments with Mer2 3A and 4A mutants. Yeast were transformed with the pGAD-C1 (activation domain) and pGBDU (DNA binding domain) plasmids as indicated. Cells were grown and pipetted onto non-selective (left) or selective plates (right) at three concentrations. B) *In vitro* pulldowns between Strep-Mre11 and Mer2. C-terminally 2xStrepII-tagged Mre11 was used to capture Mer2^WT^, Mer2^3A^ and Mer2^4A^ on Streptactin beads. Simultaneously Mer2^WT^, Mer2^3A^ and Mer2^4A^ in the absence of Mre11-Strep were used to determine non-specific binding to the beads. C) Southern blot analysis comparing double strand break (DSB) levels in *sae2*D, *mer2*D*sae2*D, *MER2-3HA mer2*D*sae2*D, *mer2-3a-3HA mer2*D*sae2*D, and *mer2-4a-3HA mer2*D*sae2*D strains. Cells were harvested at indicated time points post induction of meiosis and faster migrating DNA species (indicative of DSBs) were detected using a probe for the *YCR047C* locus. *sae2*D cells are resection-deficient and cannot repair DSBs. D) Final meiotic loop-axis cartoon. Summarises the novel interactions shown in the previous figures while highlighting the integration of Mre11via its direct binding to Mer2, into the loop axis model.

The role of Mer2 in meiotic DSB activity is key in enabling faithful meiotic chromosome segregation and viable spore formation. Consequently, *mer2*Δ strains exhibit very strong spore viability defects (^15^ and Figure 5C). As a first test of functionality, we investigated the effect of our *MER2* alleles on this defect. Expression of Mer2-3HA in *mer2*Δ overcame the spore viability defect seen in cells lacking Mer2 (Figure 5C) demonstrating functionality of our *MER2* expressing constructs. Strikingly, expression of Mer2^3A^-3HA or Mer2^4A^-3HA failed to restore spore viability in *mer2*Δ; these strains essentially behaved indistinguishably from the *mer2*Δ strain. This suggests that the designed mutants disrupt a functionality of Mer2 that is key to its role during meiotic prophase.

Since the investigated region of Mer2 is located within the N-terminal region of the protein, we expected that these mutants would not disrupt the interactions with Hop1 and nucleosomes described above. In line with this idea, we observed that *in vitro*, recombinant Mer2^3A^ and Mer2^4A^ are able to interact with nucleosomes (Figure 3F) and Hop1 (Figure 4D), albeit with slightly lower affinity for nucleosomes as determined by EMSA (Supplementary Figure 4C).

To attempt to trace the defect of these *MER2* alleles, we first investigated the association of Mer2 on spread meiotic chromosomes during meiotic prophase. We found that the recruitment of Mer2 to defined chromosomal foci was not disrupted in Mer2^3A^ and Mer2^4A^ expressing situations (Figure 5D). In contrast, quantification of the number of Mer2 foci that were observed during meiotic prophase revealed an increase in cells expressing these mutant proteins (Figure 5E). We currently do not understand the reason underlying this apparent increase in chromosome-associated Mer2 foci, but it could conceivably be related to known feedback regulation that ensures sustained DSB activity in conditions where chromosome synapsis fails or is delayed^38^. In any case, recruitment of Mer2 to chromatin-associated foci was not disrupted, suggesting that Mer2^3A^ and Mer2^4A^ are proficient to interact with upstream factors that lead to chromatin recruitment. Since HORMA proteins are suggested to drive recruitment of Mer2 to the chromosome axis (^16,17,28^ and see our earlier observations), these results also strongly argue that the defect that is triggered by *mer2-3a* and *mer2-4a* is independent of its interaction with Hop1. Furthermore, the observation that spore viability was severely impaired in cells expressing *mer2-3a* and *mer2-4a* is in contrast with the relatively mild phenotype caused by the specific disruption of the Mer2-Spp1 interaction^13^. Together, these observations, in combination with the strong evolutionary conservation of the affected region, argue that the observed defects are caused by the disruption of yet another key Mer2 interaction, potentially with another essential DSB factor.

### The Mer2 3A and 4A mutants are defective in Mre11 binding and DSB formation

To identify the underlying molecular defect underlying the observed phenotypes in our *mer2-3a* and *mer2-4a* mutants, we carried out a directed yeast-two-hybrid (Y2H) analysis against previously identified Mer2 interactors involved in DSB formation (Xrs2, Mre11, Ski8, Spo11, Rec104)^39^ and Spp1 (as a positive control). Some of the interactions were complicated by the presence of apparent self-activation by the empty vector, but we observed a clear difference in binding between the Mer2 and Mre11 when comparing wild type *MER2* to our *mer2-3a* or *mer2-4a* mutants (Figure 6A, data not shown). Importantly, these mutants were proficient to interact with Spp1, similar to what was indicated by our *in vitro* binding assays (Figure 1C).

To confirm a direct interaction between Mer2 and Mre11, we expressed and purified recombinant Mre11 carrying an Cterminal 2xStrep-II tag using baculovirus-based expression. We then used this as a bait to pulldown Mer2^WT^, Mer2^3A^ and Mer2^4A^. We observed a reduced interaction between Mre11 and both of the Mer2 mutants (Figure 6B). We note that the difference in the interaction between Mer2 and Mre11, a recombinant pulldown between Mre11 and Mer2^3A^ and Mer2^4A^ is not as striking as in the Y2H (see discussion). Since it has been previously shown that the MRX complex is generally required to make meiotic DSBs (reviewed in ^40^) including Mre11 specifically^41^, we investigated meiotic DSB formation in *mer2-3a* and *mer2-4a* mutants. We used Southern blotting to track meiotic DSB formation at *YCR047C*, a confirmed DSB hotspot. Note that we utilized the *sae2*Δ resection and repair-deficient background in order to enable DSB detection. We detected a strong impairment of meiotic DSB activity specifically in *mer2-3a* or *mer2-4a*, to an extent that was comparable to what was observed in *mer2*Δ strains (Figure 6C). As such, we propose that, in addition to its centrally positioned role in mediating chromosome axis and chromatin loop-tethering of the DSB machinery, a third key contribution of Mer2 (and its conserved N-terminal region) to DSB activity is to interact with Mre11. Presumably, not only is the presence of Mre11 required to make meiotic DSBs but its specific interaction with Mer2 is essential for the initiation of DSB formation during meiotic prophase.

## Discussion

We have biochemically dissected the function of Mer2 *in vitro* to reveal several novel features that have the potential to explain several *in vivo* characteristics of Spo11-dependent DSB formation. Firstly, we shed light on the interaction between Spp1 and Mer2. The interaction between Spp1 and Mer2 is tight (∼25 nM) and not dependent on any post-translational modification or additional cofactors. As such, the interaction between Spp1 and Mer2 is likely constitutive *in vivo*. Importantly, we also find that Mer2 serves as a dimerisation platform for Spp1, effectively increasing its affinity for H3K4^me3^ nucleosomes. Presumably this is essential given that the interaction between COMPASS bound Spp1 and H3K4^me3^ is transient, whereas the association of the DSB forming machinery is an apparently more stable event^14^. Given that the Spp1 interaction domain of Mer2 is the same as the tetramerisation domain we speculate that the antiparallel arrangement of coiled-coils of Mer2^core,19^ form two oppositely oriented binding sites for Spp1, with V195 at the centre^13,20^.

Surprisingly, we find that Mer2 itself is a bona-fide nucleosome binder. This interaction occurs at high affinity, though we assume that the true affinity is somewhat less than what we determine from EMSA titrations. Given that Mer2 has been previously shown to bind DNA^19^, yet still binds to the nucleosome core particle (with 147 bp DNA and no DNA overhangs), and that neither the loss of histone tails, nor the acidic patch mutant disrupted the interaction, we suggest that Mer2 binds to nucleosomal DNA. This idea is supported by the XL-MS data on the Spp1-Mer2 complex bound to nucleosomes, which places Mer2 proximal to the DNA entry/exit site (Figure 2D and E). Indeed the tight and specific nucleosome binding ability of Mer2 provides a molecular basis for the observation that neither Spp1, nor Set1 mediated H3K4me3 are absolutely required to make meiotic DSBs^9,10^, unlike Mer2 itself ^15^. As such we speculate that in the absence of H3K4^me3^ marks, or the Spp1 reader, Mer2 binds stochastically to nucleosomes that are positioned in chromatin loops. Such a model speculates that some of these binding events present a nucleosome depleted loop region to Spo11, but many do not, explaining also that meiotic DSBs are severely reduced in number in an *spp1* or *set1* background^9,10^. On the other hand if Mer2 preferentially bound free-DNA, as opposed to nucleosomal DNA, there might not be such a severe DSB phenotype in *spp1* or *set1* cells. The association between nucleosomes, Spp1 and Mer2 is important to establish the connection between chromosome axis-associated DSB factors and DSB sites that are localized in the loop. We speculate that the combination of ‘generic’ nucleosome binding (via Mer2-nucleosomal DNA) and specific histone tail recognition (Spp1-PHD domain-H3 tail) endows the DSB machinery with the required binding strength and combined with specificity.

Our observation, and characterisation, of the direct interaction between Hop1 and Mer2 offers a tantalising glimpse into how axial proteins may be recruiting DSB factors - through Mer2 - to regulate Spo11 activity. A key observation is that we only observed binding between Mer2 and Hop1^WT^ in the presence of Red1. We could exclude any significant direct binding to Red1 which is consistent with the observations in mammals and fission yeast, where Hop1 orthologs have been implicated as the direct interactor of Mer2 orthologs^16,17^ - though we here, for the first time, demonstrate a direct interaction *in vitro* between purified components. We propose a model whereby Hop1 that is bound to its own closure motif is not competent for Mer2 binding (Figure 4E, left), possibly due to blocking of binding interfaces present within its C-terminus. We posit that only once Hop1 is bound to Red1 the binding site for Mer2 becomes “unlocked” and accessible (Figure 4E, right). Given that one key conserved feature of meiotic HORMADs is the presence of the closure motif in the C-terminus of the protein, we propose that this is a common mechanism for the regulation of the recruitment of Mer2 orthologs to the meiotic axis.

Intriguingly, in fission yeast, the zinc-finger of Hop1 (located between residues 348-364 within the C-terminal region of *S. cerevisiae* Hop1) has been suggested to be required for Mer2 binding ^16^, which would not be inconsistent with our data. We believe that our observations provide mechanistic insights into the specific recruitment of Mer2 to meiotic chromosomes. For example, this model might explain how Mer2 can specifically be recruited to meiotic chromosomes, and not form spurious interactions with non-chromosomal Hop1: non-chromosomal, monomeric Hop1 is thought to be largely present in the intramolecular ‘closed’ form (unless it is converted into the ‘open/unbuckled’ state by Pch2/TRIP13, which is thought to promote rapid chromosomal incorporation of Hop1)^30,42^. Conversely, it might explain how DSB activity is negatively regulated by chromosome synapsis. Synapsis leads to recruitment of Pch2/TRIP13, and removal of Hop1 from the chromosomal axis (presumably by opening-up Hop1 bound to Red1). This would be associated with immediate co-removal of Mer2, and once released Hop1 would transition into intramolecular Hop1, leading to disruption of the Mer2-Hop1 complex ^29,30,31,33,34,43^.

Finally, we attempted to create separation of function mutants for Mer2 by mutating a previously undescribed conserved region in the N-terminus of the protein. Both mutants in this region (3A and 4A) were very penetrative and exhibited a complete loss of spore viability (Figure 5C). Nonetheless the mutants still showed normal localisation to the meiotic axis *in vivo*, and maintained an interaction with Hop1 *in vitro*. Interestingly, although quantitative EMSAs showed that there was a reduction in affinity of Mer2^3A^ for nucleosomes (Supplementary figure 4C) but under physiological salt conditions (*i*.*e*. in the pulldown in Figure 3F) both Mer2 mutants could still form a complex with nucleosomes.

Interestingly, we discovered that these mutants interfere with an interaction between Mer2 and Mre11. We note that the observed severity of the effect of the Mer2^3A^ and Mer2^4A^ on the Mer2-Mre11 interaction: yeast-2-hybrid analysis indicated a strong disruption of the interaction when using Mer2^3A^ or Mer2^4A^ compared to Mer2. Interestingly, while we also observed interaction between Mer2 and Mre11 using *in vitro* purified proteins, and that the mutants reduced the interaction between Mer2 and Mre11, the difference between the mutants and wild-type was less pronounced. We infer that a factor/condition that strengthens the interaction between Mer2 and Mre11 is present in vegetatively growing yeast cells (*i*.*e*. the condition of Y2H analysis), but is lacking in our *in vitro* pulldown. This could be an additional protein factor, perhaps one of the other components of the Mre11-Rad50-Xrs2 complex, or a post-translational modification. For example, Mer2 has been previously shown to be phosphorylated by DDK and S-Cdk^8,44^, and Mer2^3A^ or Mer2^4A^ could potentially affect these phosphorylation events. Moreover, Mre11 has been previously shown to contain two SUMO interaction motifs (SIMs). Since many meiotic proteins have been recently shown to be SUMOylated in yeast, among them Mer2, SUMOylation might serve as an important regulator^45^. Importantly the residues of the Mer2-3A mutants are essentially universally conserved throughout eukaryotes (Supplementary Figure 6) and as such we would expect that the direct interaction between Mer2 and Mre11, via these residues, is a universal feature of the meiotic program.

Taken together our data show that Mer2 forms the keystone of meiotic recombination, binding directly to both the axis via Hop1 and the loop via nucleosomes. Presumably once assembled on the loop-axis Mer2 is then able to interact with Rec114 and Mei4 in phospho-dependent manner (likely via the PH domains of Rec114^28,46,47^), an interaction that may also be somehow further controlled by liquid-liquid phase separation^20^. Simultaneously the N-terminus of Mer2 presumably recruits the MRX complex via Mre11, a step that is critical for the formation of meiotic DSBs and the initiation of meiosis. In organisms with a synaptonemal complex (SC), shutting meiotic DSB formation is associated with synapsis. It has been shown that synapsis results in the Pch2 mediated displacement of Hop1 from the axistextsuperscript34,48,49). We expect that this action would also result in the displacement of Mer2 as Hop1 would “snap shut” and further interaction with Hop1 would be prevented via steric hindrance of the binding site.

Our findings reveal the power of a biochemical reconstitution in dissecting the function of complex biological systems. Mer2 emerges as the keystone of meiotic recombination, interacting with the axis, nucleosomes, the DSB forming machinery, and the repair machinery. Larger and more ambitious reconstitutions would enable us to probe the role of additional protein cofactors and posttranslational modifications in meiotic regulation.

## Materials and Methods

### Protein expression and Purification

Sequences of *Sac-charomyces cerevisiae SPP1, HOP1, RED1* and *MRE11* were derived from SK1 strain genomic DNA. Due to the presence of an intron in *MER2*, this was amplified as two separate fragments and Gibson assembled together.

All Mer2 constructs were expressed as an 3C HRV cleavable N-terminal MBP fusion in chemically competent C41 *E. coli* cells. Protein expression was induced by addition of 250 µM IPTG and the expression continued at 18°C overnight. Cells were washed with 1xPBS and resuspended in lysis buffer (50 mM HEPES pH 7.5, 300 mM NaCl, 5 % glycerol, 0.1 % Triton-X 100, 1 mM MgCl2, 5 mM beta-mercaptoethanol). Resuspended cells were lysed using an EmulsiFlex C3 (Avestin) in presence of DNAse (10 µg/ml) and AEBSF (25 µg/ml) before clearance at 20,000g at 4C for 30 min. Cleared lysate was applied on a 5 mL MBP-trap column (GE Healthcare) and extensively washed with lysis buffer. Mer2 constructs were eluted with a lysis buffer containing 1 mM maltose and passed through a 6 mL ResourceQ column (GE Healthcare) equilibrated in 50 mM HEPES pH 7.5, 100 mM NaCl, 5% glycerol, 5 mM betamercaptoethanol. The proteins were eluted by increasing salt gradient to 1M NaCl. Protein containing elution fractions were concentrated on Amicon concentrator (100 kDa MWCO) and loaded a Superose 6 16/600 (GE Healthcare) pre-equilibrated in SEC buffer (50 mM HEPES pH 7.5, 300 mM NaCl, 10% glycerol, 1 mM TCEP). Untagged Mer2^FL^ was prepared likewise until concentration of protein eluted from ResourceQ. The concentrated eluent was mixed with 3C HRV protease in a molar ratio of 50:1 and incubated at 4C for 6 hours. Afterwards, the cleaved protein was loaded on a Superose 6 16/600 pre-equilibrated in SEC buffer for cleaved Mer2 (20 mM HEPES pH 7.5, 500 mM NaCl, 10% glycerol, 1 mM TCEP, 1 mM EDTA, AEBSF).

Spp1 constructs were produced as an 3C HRV cleavable N-terminal MBP or GST fusion in a similar manner as MBP-Mer2. To purify GST-Spp1, cleared lysate was applied on GST-Trap (GE Healthcare) before extensive washing with lysis buffer. The protein was eluted with a lysis buffer with 40 mM reduced glutathione and passed through ResourceQ. Both GST and MBP could be cleaved by adding 3C HRV protease to concentrated protein (using an Amicon concentrator with 30 kDa cutoff) in 1:50 molar ratio. After a ∼6 hour incubation at 4°C, the cleaved protein was loaded on Superdex 200 16/600 pre-equilibrated in SEC buffer (50 mM HEPES pH 7.5, 300 mM NaCl, 10% glycerol, 1 mM TCEP).

Hop1 constructs were produced as 3C HRV cleavable N-terminal Twin-StrepII tag in BL21 STAR *E*.*coli* cells. The expression was induced by addition of 250 µM IPTG and the expression continued at 18°C for 16 hours. Cleared lysate was applied on Strep-Tactin Superflow Cartridge (Qiagen) before extensive washing in lysis buffer. The bound protein was eluted with a lysis buffer containing 2.5 mM desthiobiotin and loaded on HiTrap Heparin HP column (GE Health-care) and subsequently eluted with increasing salt gradient to 1M NaCl. Eluted Strep-Hop1 constructs were concentrated on a 30 kDa MWCO Amicon concentrator and loaded on Superdex 200 16/600 pre-equilibrated in SEC buffer.

Red1 was produced in insect cells as a C-terminal MBP-fusion either alone or co-expressed with Strep-Hop1. Bacmids were in both cases produced in EmBacY cells and subsequently used to transfect Sf9 cells to produce baculovirus. Amplified baculovirus was used to infect Sf9 cells in 1:100 dilution prior to 72 hour cultivation and harvest. Cells were extensively washed and resuspended in Red1 lysis buffer (50 mM HEPES pH 7.5, 300 mM NaCl, 10% glycerol, 1 mM MgCl2, 5 mM beta-mercaptoethanol, 0.1% Triton-100). Resuspended cells were lysed by sonication in presence of Benzonase and a protease inhibitor cocktail (Serva) before clearance at 40,000g at 4C for 1h. Cleared lysate was loaded on Strep-Tactin Superflow Cartridge (in case of Red1-Hop1 complex) or MBP-trap column (in case of Red1 alone). Proteins were eluted using a lysis buffer containing 2.5 mM desthiobiotin and 1 mM maltose, respectively. Partially purified proteins were further passed through HiTrap Heparin HP column and eluted with increasing salt gradient to 1M NaCl. Purified proteins were subsequently concentrated using Pierce concentrator with 30 kDa cutoff in 50 mM HEPES pH 7.5, 300 mM NaCl, 10% glycerol, 1 mM TCEP. Because of the small yield of the proteins, the SEC purification step was neglected and the purity of the proteins was checked using the Refeyn One mass photometer.

Mre11 was produced as a C-terminal Twin-StrepII tag in insect cells using the same expression conditions as for Red1 protein. The cell pellet was resuspended in Mre11 lysis buffer (50 mM HEPES pH 7.5, 300 mM NaCl, 5% glycerol, 0.01% NP40, 5 mM β-mercaptoethanol, AEBSF, Serva inhibitors). Resuspended cells were lysed by sonication before clearance at 40,000g at 4C for 1h. Cleared lysate was loaded on a 5-ml Strep-Tactin XT Superflow Cartridge (IBA) followed by first wash using 25 ml of high-salt wash buffer (20 mM HEPES pH 7.5, 500 mM NaCl, 5% glycerol, 0.01% NP40, 1 mM β-mercaptoethanol) and second wash step using 25 ml of low-salt wash buffer (20 mM HEPES pH 7.5, 150 mM NaCl, 5% glycerol, 0.01% NP40, 1 mM β-mercaptoethanol). The Mre11 protein was eluted with 50 mL of low-salt wash buffer containing 50 mM biotin. Partially purified protein was further loaded onto a 5-ml Heparin column (GE Healthcare) pre-equilibrated in a low-salt wash buffer and eluted with increasing salt gradient to 1 M NaCl. The fractions containing Mre11 protein were concentrated on a 50 kDa MWCO Amicon concentrator and applied onto a Superdex 200 10/300 column (GE Healthcare) pre-equilibrated in Mre11 SEC buffer (20 mM HEPES pH 7.5, 300 mM NaCl, 5% glycerol, 1 mM β-mercaptoethanol, 1 mM TCEP).

### SEC-MALS analysis

50 µL samples at 5-10 µM concentration were loaded onto a Superose 6 5/150 analytical size exclusion column (GE Healthcare) equilibrated in buffer containing 50 mM HEPES pH 7.5, 1 mM TCEP, 300 mM NaCl (for samples without nucleosomes) or 150 mM NaCl (for samples with nucleosomes) attached to an 1260 Infinity II LC System (Agilent). MALS was carried out using a Wyatt DAWN detector attached in line with the size exclusion column.

### Microscale thermophoresis

Triplicates of MST analysis were performed in 50 mM HEPES pH 7.5, 300 mM NaCl, 5% glycerol, 1 mM TCEP, 0.005% Tween-20 at 20°C. The final reaction included 20 uM RED-NHS labelled untagged Spp1 (labelling was performed as in manufacturer’s protocol-Nanotemper) and titration series of MBP-Mer2 constructs (concentrations calculated based on oligomerization stage of Mer2). The final curves were automatically fitted in Nanotemper analysis software.

### Recombinant Nucleosome production

Recombinant *X. laevis* histones were purchased from “The Histone Source” (Colorado State) with the exception of H3-C110A_K4C cloned into pET3, which was kindly gifted by Francesca Matirolli. The trimethylated H3 in C110A background was prepared as previously described^21^. *X. laevis* histone expression, purification, octamer refolding and mononucleosome reconstitution were performed as described^50^. Plasmids for the production of 601-147 (pUC19) and 601-167 (pUC18) DNA were kindly gifted by Francesca Matirolli (Hubrecht Institute, Utrecht) and Andrea Musacchio (MPI Dortmund), respectively. DNA production was performed as previously described ^50^. Reconstituted mononucleosomes were shifted to 20 mM Tris pH 7.5, 150 mM NaCl, 1 mM EDTA, 1 mM TCEP with addition of 20% glycerol prior to freezing in - 80C.

### Electrophoretic mobility shift assays

Quadruplate EM-SAs were carried out as previously described^51^, at a constant nucleosome/NCP/DNA concentration of 10 nM with the DNA being post-stained with SybrGold (Invitrogen). Gels were imaged using a ChemiDocMP (Bio-Rad Inc). Nucleosome depletion in each lane was quantitated byImageJ, using measurements of triplicate of the nucleosome alone for each individual gel as a baseline. Binding curves were fitted using Prism software and the following algorithm (Y=Bmax*X^h/(Kd^h + X^h)). It was necessary in each Mer2 case to add a Hill coefficient to obtain the best fit.

### Streptactin pulldown

Streptactin pulldowns were performed using pre-blocked Streptavidin magnetic beads (Pierce) in a pulldown buffer (20 mM HEPES pH 7.5, 300 mM NaCl, 5% glycerol, 1 mM TCEP). 1 uM Strep-Hop1 as a bait was incubated with 3 µM Mer2 as a prey in 40 µL reaction for 2 hours on ice without beads and another 30 min after addition of 10 µL of beads pre-blocked with 1 mg/ml BSA for 2 hours. After incubation, the beads were washed twice with 200 µL of buffer before elution of the proteins with a 1x Laemmli buffer. Samples were loaded on 10% SDS-PAGE gel and afterwards stained with InstantBlue.

### Amylose pulldown

Amylose pulldowns were performed using pre-blocked Amylose beads (New England BioLabs) in a pulldown buffer (20 mM HEPES pH 7.5, 300 mM NaCl, 5% glycerol, 1 mM TCEP). 1 uM Red^I743R^-MBP or Red^I743R^-MBP/Strep-Hop1 as a bait was incubated with 3 µM Mer2 as a prey in 40 µL reaction for 2 hours on ice without beads and another 1 hour after addition of 10 µL of beads pre-blocked with 1 mg/ml BSA for 2 hours. After incubation, the beads were washed twice with 200 µL of buffer before elution of the proteins with a buffer containing 1 mM maltose. Samples were loaded on 10% SDS-PAGE gel and afterwards stained with InstantBlue.

### Biotinylated nucleosome pulldown

Biotinylated nucleo-somes (0.5 µM) or NCP (0.4 µM) were incubated with prey proteins (1.5 µM) for 30 min on ice in buffer containing 20 mM HEPES pH 7.5, 150 mM NaCl, 5% glycerol, 1 mM EDTA, 0.05% Triton-X100, 1 mM TCEP in a reaction volume of 40 µL. 10 µL of protein mix were taken as an input before adding 10 µL of pre-equilibrated magnetic Dynabeads M 270 streptavidin beads (Thermo Fisher Scientific) to the reaction. The samples with beads were incubated on ice for 2 min before applying magnet and removing the supernatant. The beads were washed twice with 200 µl of buffer. To release the streptavidin from the beads, Laemmli buffer (1x) was added to the beads and incubated for 10 min. Samples were analyzed on 10-20% SDS-PAGE gel and stained by InstantBlue.

### Analytical Size exclusion chromatography

Analytical SEC was performed using Superose 6 5/150 GL column (GE Healthcare) in a buffer containing 20 mM HEPES pH 7.5, 150 mM NaCl, 5% glycerol, 1 mM TCEP, 1 mM EDTA. All samples were eluted under isocratic elution at a flow rate of 0.15 ml/min. Protein elution was monitored at 280 nm. Fractions were subsequently analysed by SDS-PAGE and Instant-Blue staining. To detect complex formation, proteins were mixed at 5 uM concentration in 50 µL and incubated on ice for 1 hour prior to SEC analysis.

### Mass Photometry

Mass Photometry was performed in 20 mM HEPES pH 7.5, 150 mM NaCl, 5% glycerol, 1 mM TCEP, 1 mM EDTA. Mer2 and mononucleosomes (600 nM) were mixed and incubated for 1 hour on ice prior to analysis using the Refeyn One mass photometer. Immediately before analysis, the sample was diluted 1:10 with the aforementioned buffer. Molecular mass was determined in Analysis software provided by the manufacturer using a NativeMark (Invitrogen) based standard curve created under the identical buffer composition.

### Cross-linking Mass Spectrometry (XL-MS)

For XL-MS analysis proteins were dissolved in 200 ul of 30 mM HEPES pH 7.5, 1 mM TCEP, 300 mM NaCl (for samples without nucleosomes) or 150 mM NaCl (for samples with nucleosomes) to final concentration of 3 µM, mixed with 3 µL of DSBU (200 mM) and incubated for 1 hour in 25 °C. The reaction was quenched by addition of 20 µL of Tris pH 8.0 (1 M) and incubated for another 30 min in 25 °C. The crosslinked sample was precipitated by addition of 4X volumes of 100% cold acetone ON in -20 °C and subsequently analysed as previously described^52^.

### Yeast strains

All strains, except those used for yeast two-hybrid analysis, are of the SK1 background. See Supplementary Material for a description of genotypes of strains used per experiment. For MER2 alleles, constructs containing pMER2(−1000 to 1), the coding sequence of *MER2* lacking its intron *(i*.*e*. wildtype, 3a or 4a sequences), C-terminal 3HA tag, and 500 base pair of downstream sequence flanked by HindIII and Nar1 restriction enzymes were cus- tom synthesized by Genewiz Inc. These constructs were recloned in a YIPlac128 plasmid carrying *LEU2* using restriction cloning. These plasmids were integrated at *pMER2* in front of *mer2::HISMX6* following EcoRI linearization. Correct single copy integration was confirmed by PCR.

### Growth conditions for synchronous meiosis

Yeast strains were patched onto YP-Glycerol plates and transferred to YP-4%Dextrose plates. After this, cells were grown overnight in liquid YPD culture (room temperature) followed by inoculation in pre-sporulation media (BYTA; 50 mM Sodium Phthalate-buffered, 1% yeast extract, 2% tryptone and 1% acetate) at OD600 = 0.3. Cells were grown for 18 hours in BYTA at 30°C, washed twice in water and re-suspended in sporulation media (0.3% potassium acetate) at OD600 = 1.9 to induce meiosis at 30°C.

### Spore viability and efficiency

Cells were synchronously induced into meiosis, and incubated for 24 hours. Of each strain, 200 cells were counted using a standard bright field microscope, and monads, dyads and tetrads were scored. For viability, the indicated number of tetrads was dissected using standard manipulation methods, and grown on YPD plates. Spore viability was calculated as a percentage of the total number of viable spores.

### Surface spreading of chromosomes and immunofluo-rescence

2 mL of meiotic cells (t=3 hours induction) were collected at indicated time points, killed by addition of 1% sodium azide and processed. Cells were treated with 200mM Tris pH 7.5, 20mM dithiothreitol (DTT) for 2min at room temperature followed by spheroplasting at 30°C in 1M sorbitol, 2% potassium acetate, 0.13 µg/µL zymolyase (20 minutes). Spheroplasts were washed two times in 1 mL ice-cold MES-Sorbitol solution (1M sorbitol, 0.1 M MES pH 6.4, 1mM EDTA, 0.5mM MgCl2) and resuspended in 55 µL of MES-Sorbitol. 20 µL of spheroplasts were placed on clean glass slides (that were dipped in ethanol overnight and air-dried) and 2X volumes of fixing solution (3% paraformaldehyde, 3.4% sucrose) was added. This was followed by addition of four volumes of 1% Lipsol, and mixing through gentle rotation. After one minute, 4X volumes of fixing solution were added. A glass rod was used to mechanically spread chromosomes, after which samples were dried overnight at room temperature, and stored at 20°C. Slides were treated with 0.4% Photoflo (Kodak)/PBS for 3 minutes, after which slides were dipped in PBS with gentle shaking (5 minutes). Samples were blocked by incubation with 5% BSA in PBS for 15 minutes at room temperature. Overnight incubation with desired primary antibodies was performed in a humidified chamber at 4 °C, after which slides were subjected to 2X washes of 10 minutes in PBS with gentle shaking followed by incubation with fluorescent-conjugated secondary antibody for 3 hours at room temperature. The slides were washed 2X and mounted using 20µl of Vectashield mounting media containing 4’,6-Diamidine-2’-phenylindole dihydrochloride (DAPI) (Vector Laboratories). Chromosome surface spreads were immunostained with rat *–*-HA at 1:200 (Roche) and rabbit *–*-Hop1 at 1:200 (home made)). Hop1 production was performed at the antibody facility of the Max-Planck-Institute of Molecular Cell Biology and Genetics (Dresden, Germany) using affinity purified full length 6xHis-tagged Hop1. Secondary antibodies were used at the following concentrations: Alexa 488-conjugated donkey anti-rat at 1:500; Texas Red 594-conjugated donkey anti-rabbit at 1:500. Image acquisition was done by obtaining serial z-stacks of 0.2 µm thickness at room temperature using 100x 1.42 NA PlanApo-N objective (Olympus) on a DeltaVision imaging system (GE Healthcare) equipped with an sCMOS camera (PCO Edge 5.5). The z-stack images were deconvolved using SoftWoRx software. Quantifications for the number of foci of Mer2-HA, Mer2-3A-HA Mer2-4A-HA were done using the ‘Spots’ function of the Imaris software (Bitplane). Prism 8 (GraphPad) was used to generate the Scatter plots. Statistical significance was assessed by performing Mann-Whitney U-test. For representative images, Fiji/ImageJ software was used to obtain maximum intensity projection images.

### Western blot analysis

For Western blot analysis, protein lysates from yeast meiotic cultures were prepared using trichloroacetic acid (TCA)-precipitation and run on 8 or 10% SDS-gels, transferred for 90 minutes at 300 mA and blotted with the selected antibodies, as described^53^. Primary antibodies with respective dilutions were used: rabbit *–*-Hop1 (made in-house; 1:10000), rabbit *–*-Mer2^40-271^ (made in-house; 1:10000); mouse *–*-Pgk1 (Thermo Fisher, 1:5000); rabbit *–*-phospho-Histone-H3-Thr11 (Abcam, 1:1000), mouse *–*-HA (Biolegend, diluted 1:500).

### Southern blot analysis

For Southern blot assay, DNA from meiotic samples was prepared as described^54^. DNA was digested with HindIII (to detect DSBs at the control *YCR047C* hotspot) followed by gel electrophoresis, blotting of the membranes and radioactive (32P) hybridization using a probe for *YCR047C* (chromosome *III*; 209,361–201,030)^30^. DSBs signals were monitored by exposure of an X-ray film which was analyzed using a Typhoon Trio scanner (GE Healthcare) after one week.

### Yeast two hybrid

*MRE11, SPP1* and *MER2* variants were cloned into pGAD-C1 or pGBDU-C1 vectors, respectively. The resulting plasmids were co-transformed into the *S. cerevisiae* reporter strain (yWL365) and plated onto the selective medium lacking leucine and uracil. For drop assay, 2.5 µL from 10-fold serial dilutions of cell cultures with the initial optical density (OD_600_) of 0.5 were spotted onto -Leu -Ura (control) and -Leu -Ura -His plates. Cells were grown at 30°C for several days.

## Acknowledgements

We are extremely grateful to Francesca Mattiroli (Hubrecht Institute, Utrecht), for assistance with producing mononucleosomes. The histone H3 expression plasmid pET3 and the plasmid pUC19_601-147 DNA were a kind gift from Francesca Mattiroli. The plasmid pUC18_601-167 DNA was a kind gift from Andrea Musacchio (MPI of Molecular Physiology, Dortmund). We thank Christopher Heim (MPI Developmental Biology) for technical help with SEC-MALS and MST measurements. We thank Andreas Blaha and Constanze Gremmelmaier for their technical support. Work in the Weir lab is funded by the Max Planck Society and DFG grant WE 6513/2-1. Work in the Vader lab was funded by Max Planck Society and the European Research Council (ERC StG URDNA; agreement no. 638197). SKF is funded by a studentship from the International Max Planck Research School (IMPRS) “From Molecules to Organisms”.

## Author Contributions

Cloning, expression and purification: DR, VA, SKF, DL, HR, JRW; Recombinant nucleosome production: DR; Biochemical and biophysical assays: DR; Yeast two-hybrid assays: VA; Yeast genetics, molecular cell biology and microscopy: VN, VBR and GV.; Mass Spectrometry preparation: FM.; Supervision of yeast experiments: GV. Data analysis: DR, VN, VA, PJ, GV, JRW; Supervision: JRW, GV; Drafting of manuscript: GV, JRW, DR; Funding acquisition: JRW and GV.

## Supplementary Data

Yeast strains used in this study

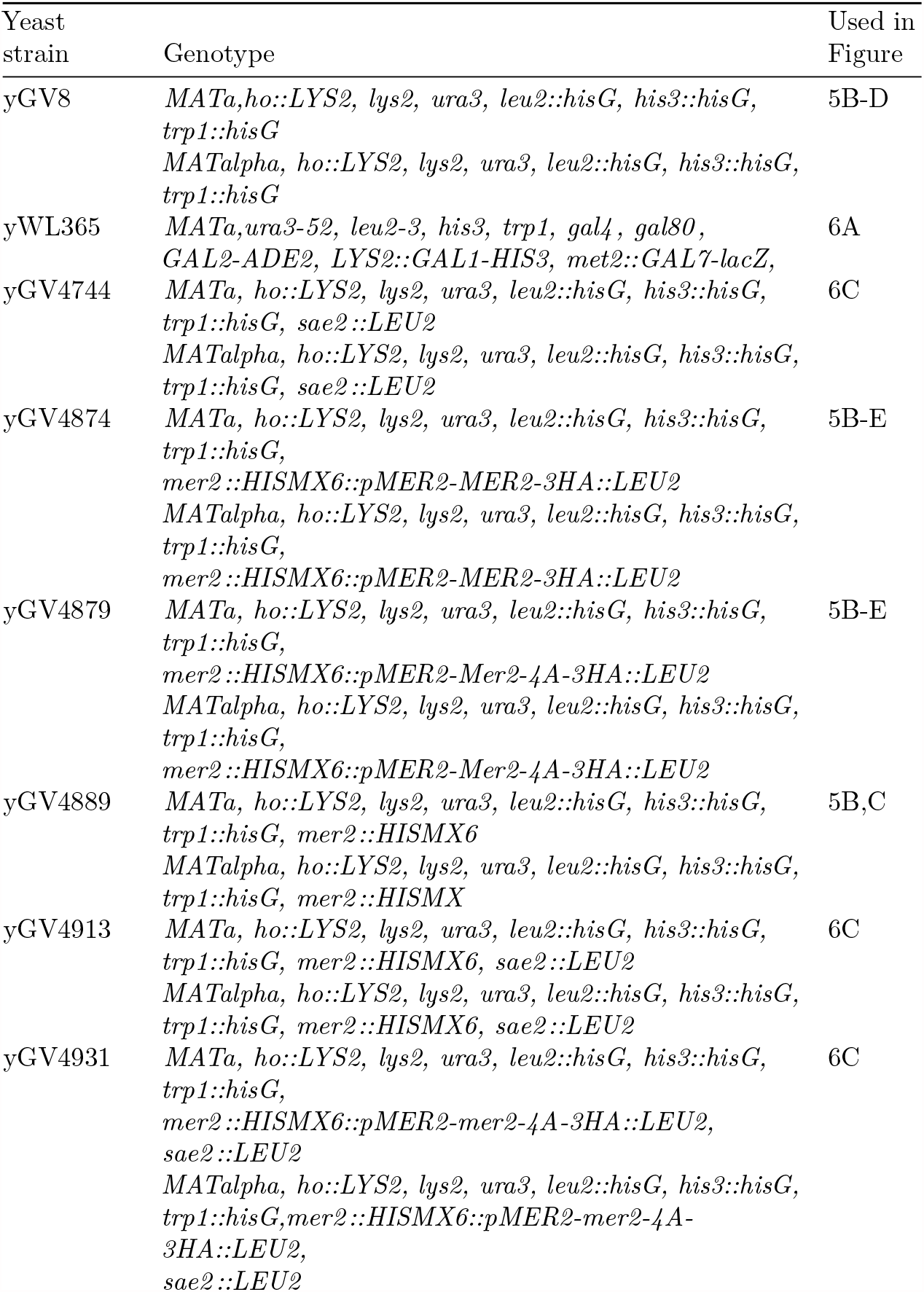
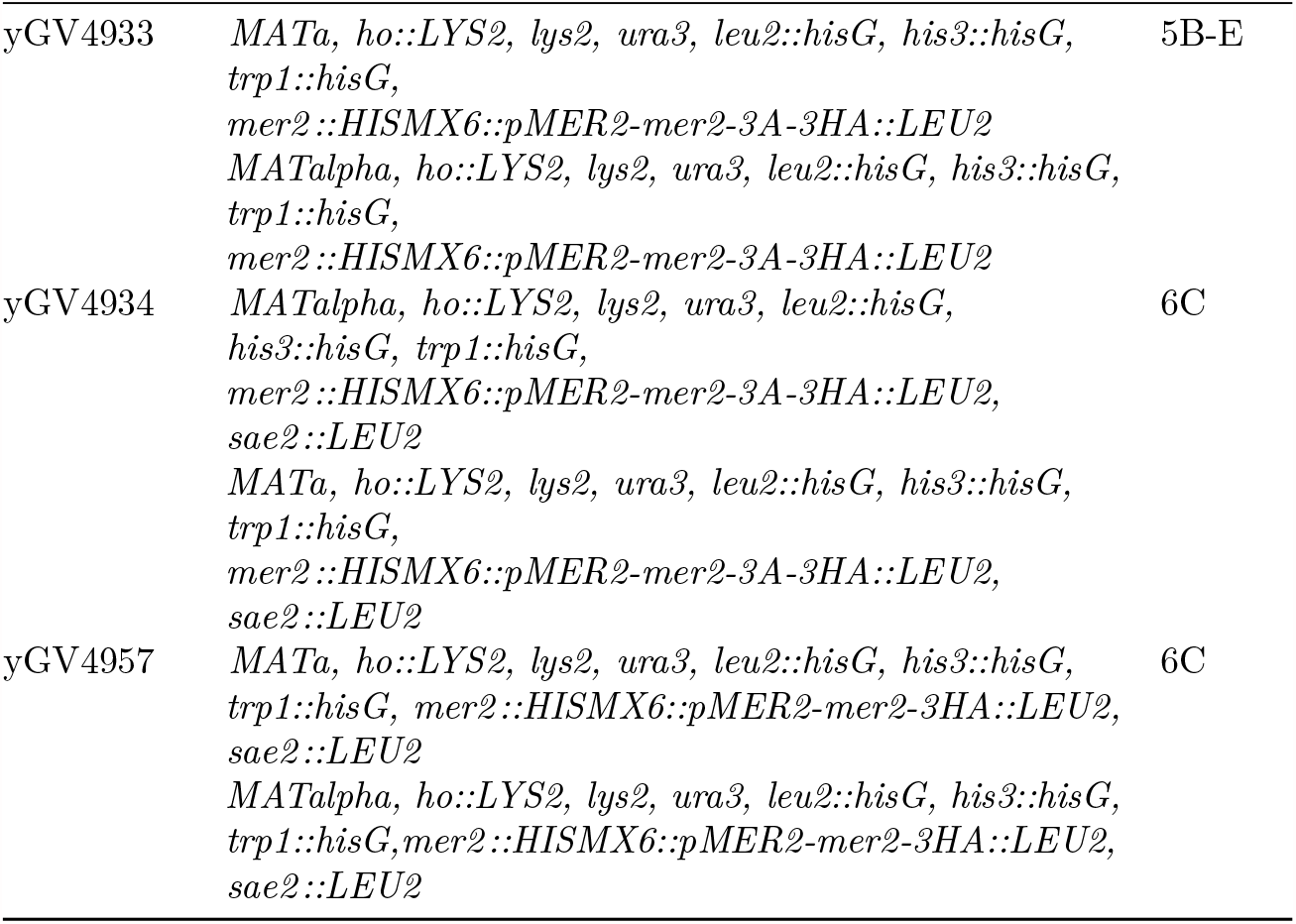

**Supplementary Figure 1.**
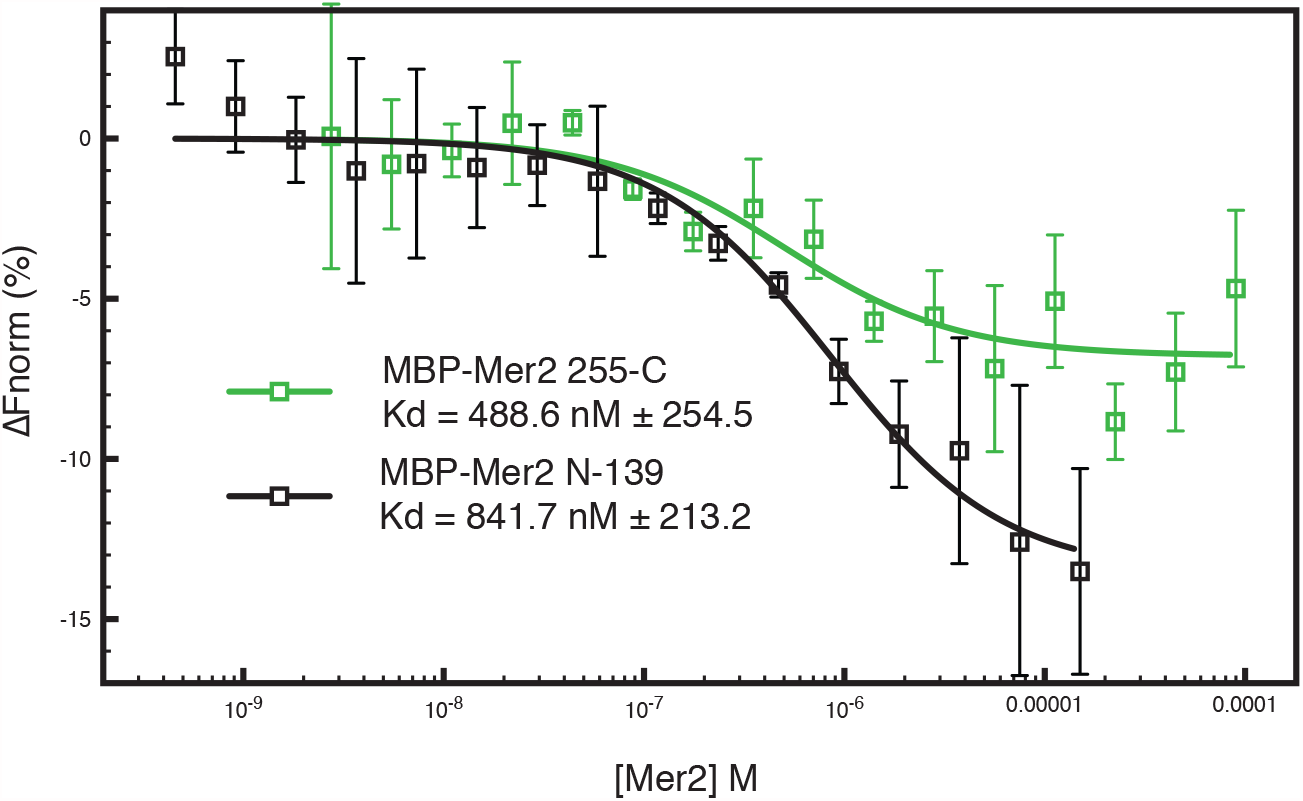
MST Measurements on additional Mer2 constructs Two MBP-tagged Mer2 constructs, MBP-Mer2 255-C (green trace) and MBP-Mer2 N-139 (black trace) were titrated against labelled Spp1 (as in Figure 1). Error bars are the SD from

**Supplementary Figure 2.**
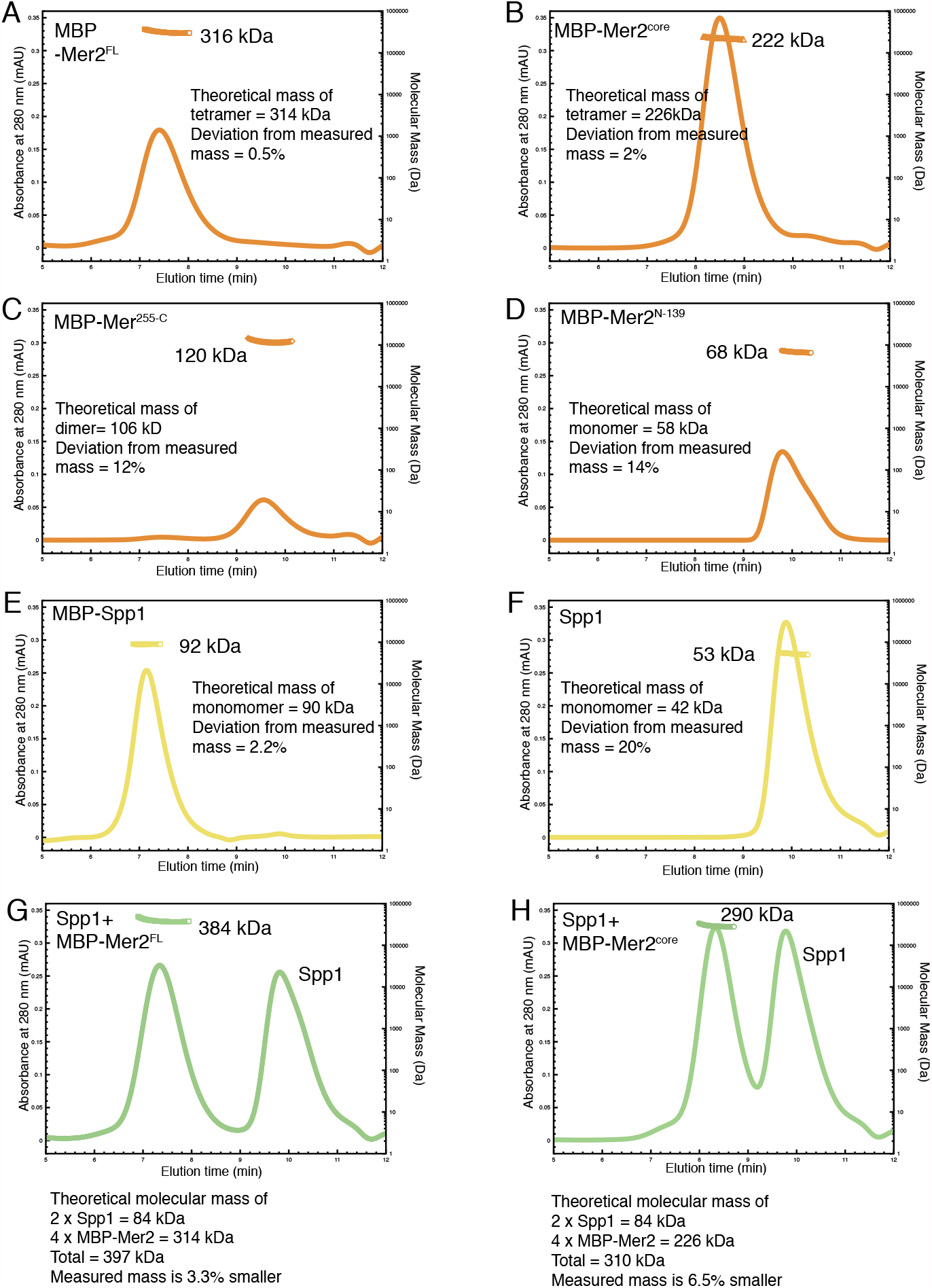
Additional SEC-MALS chromatographs and summary SEC-MALS experiments were carried out as described in materials and methods. A) N-terminally tagged MBP-Mer2^FL^ B) N-terminally tagged MBP-Mer2^core^ C) N-terminally tagged MBP-Mer2 255-C; D) N-terminally tagged MBP-Mer2 1-139; E) N-terminally tagged MBP-Spp1; F) Untagged Spp1; G) A mixture of untagged Spp1 with N-terminally tagged MBP-Mer2^FL^ the excess of Spp1 is in the right hand peak; H) A mixture of untagged Spp1 with N-terminally tagged MBP-Mer2^core^ the excess of Spp1 is in the right hand peak.

**Supplementary Figure 3.**
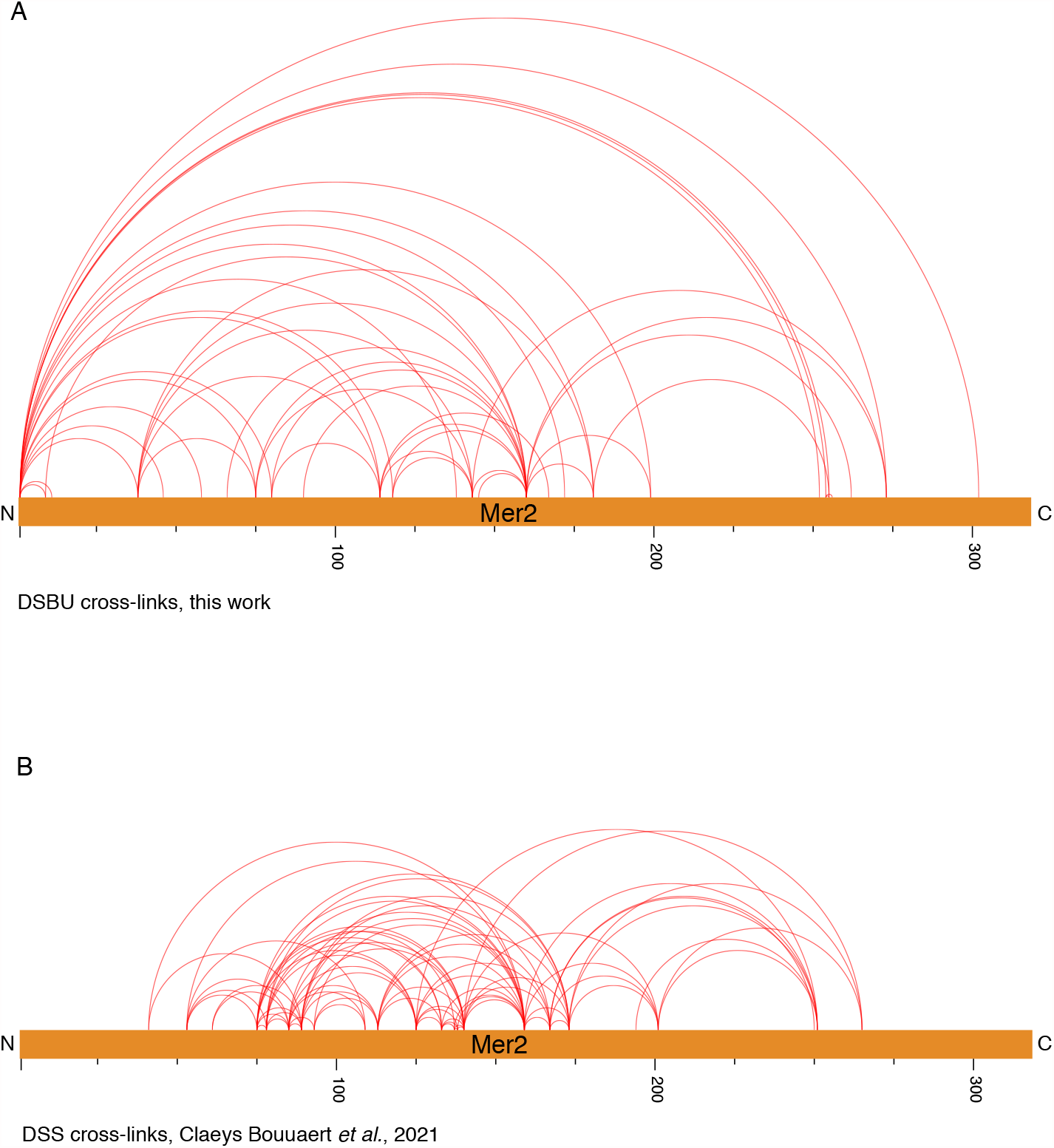
XL-MS on Mer2 alone A) Visualisation of DSBU cross-linked Mer2^FL^ alone B) Visualisation of the DSS crosslinked Mer2 sample from Bouuaert et al.^19^. Both images prepared using XVis^55^

**Supplementary Figure 4.**
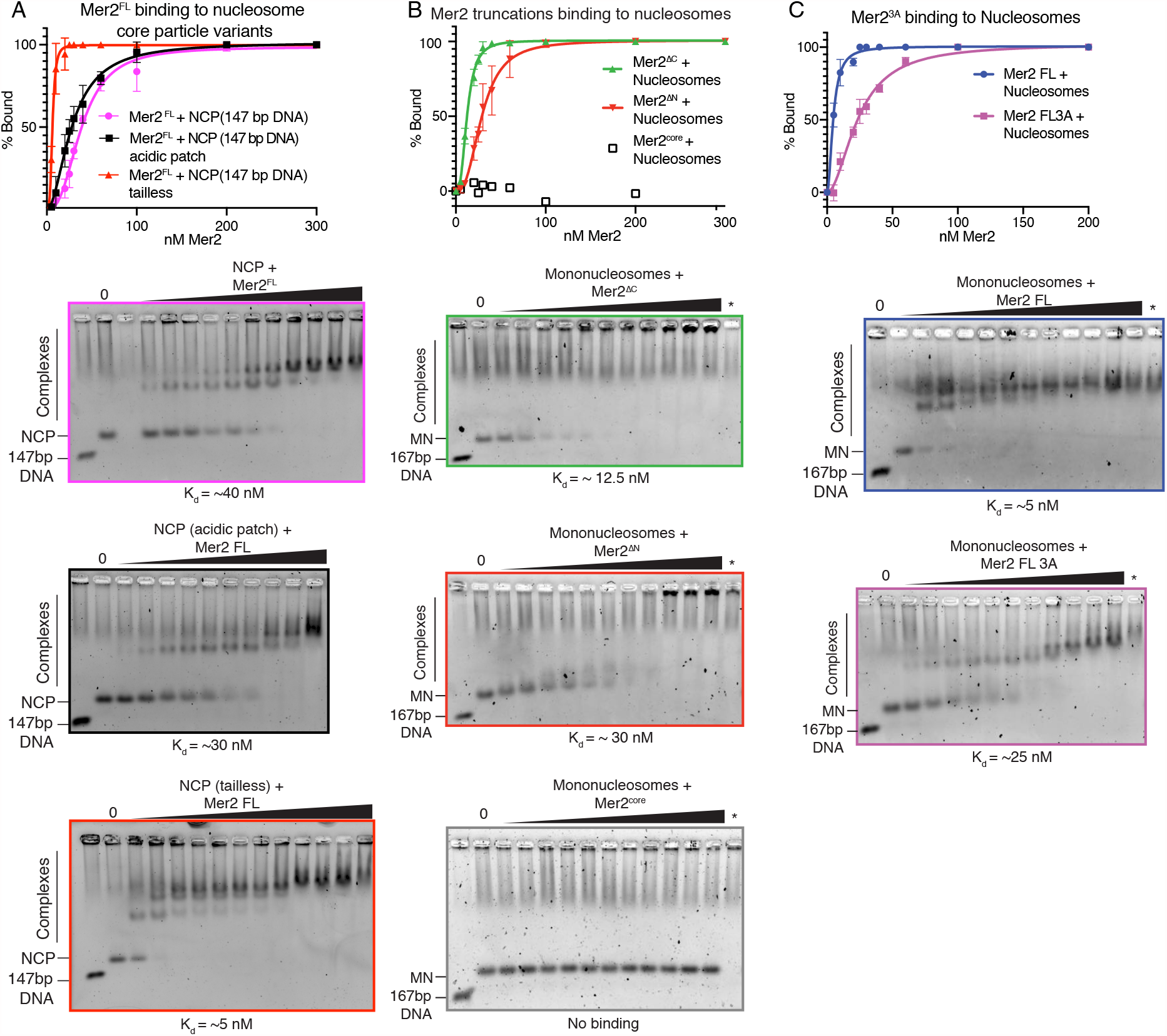
EMSAs on Mer2 binding to nucleosomes A) Mer2 binding to nucleosome core particle (NCP) variants. Mer2^FL^ was titrated against different variants of the NCP. Either WT NCPs (pink), acidic patch (56T-E61T-E64T-D90S-E91T-E92T) NCPs (black) or tailless NCPs (red) were used at a constant concentration of 10 nM. Error bars indicate the SD of three independent experiments. B) Mer2 truncation binding to mononucleosomes. Different Mer2 constructs were titrated against mononucleosomes at a constant concentration of 10 nM. Mer2^C^ (green), Mer2^N^ (red) or Mer2^core^ were used. Error bars indicate the SD of three independent experiments. C) Mer2^3A^ binding to mononucleosomes. Either Mer2^FL^ (blue trace) or Mer2^3A^ (purple trace) titrated against mononucleosomes. Error bars indicate the SD of three independent experiments.

**Supplementary Figure 5.**
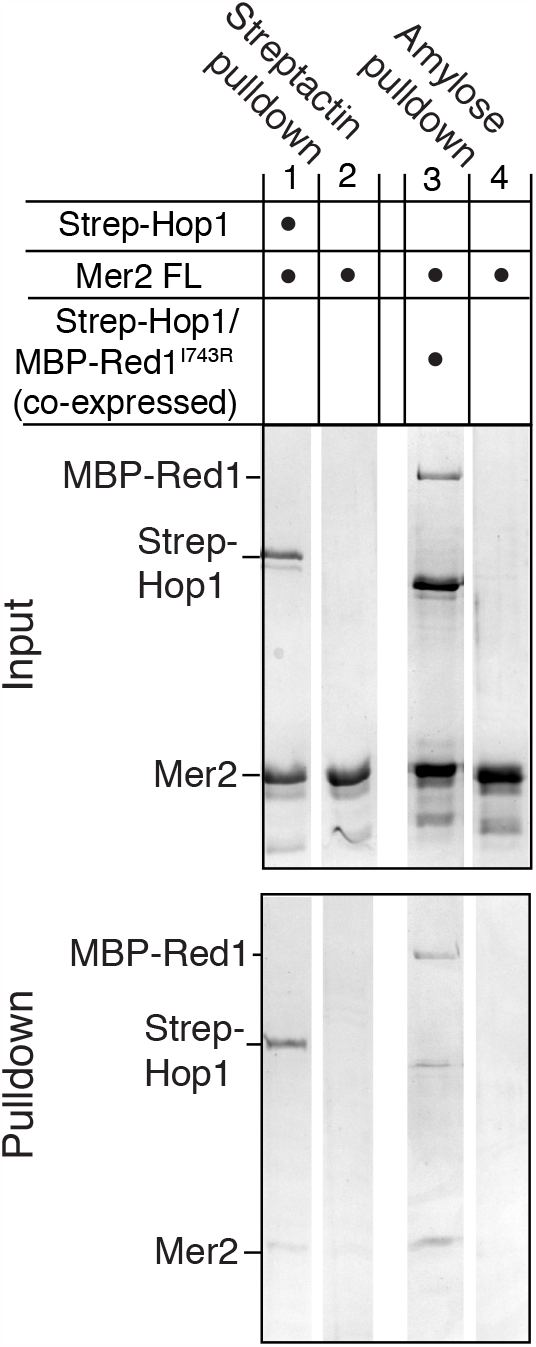
Mer2 pulldowns on Hop1-Red1 complexes Mer2^FL^ was incubated either with, or without N-terminally tagged 2xStrep-II-Hop1. Samples were then incubated with Streptactin beads to either capture the complex (lane 1), or to determine background binding of Mer2 (lane 2). Mer2^FL^ was also incubated with, or without a co-expressed and purified complex of 2xStrep-II-Hop1 and Red1-I743R-MBP. To exclude the excess Hop1 in the complex, amylose beads were used to pull down Red1 (ane 3) or to determine background binding of Mer2 (lane 4).

**Supplementary Figure 6.**
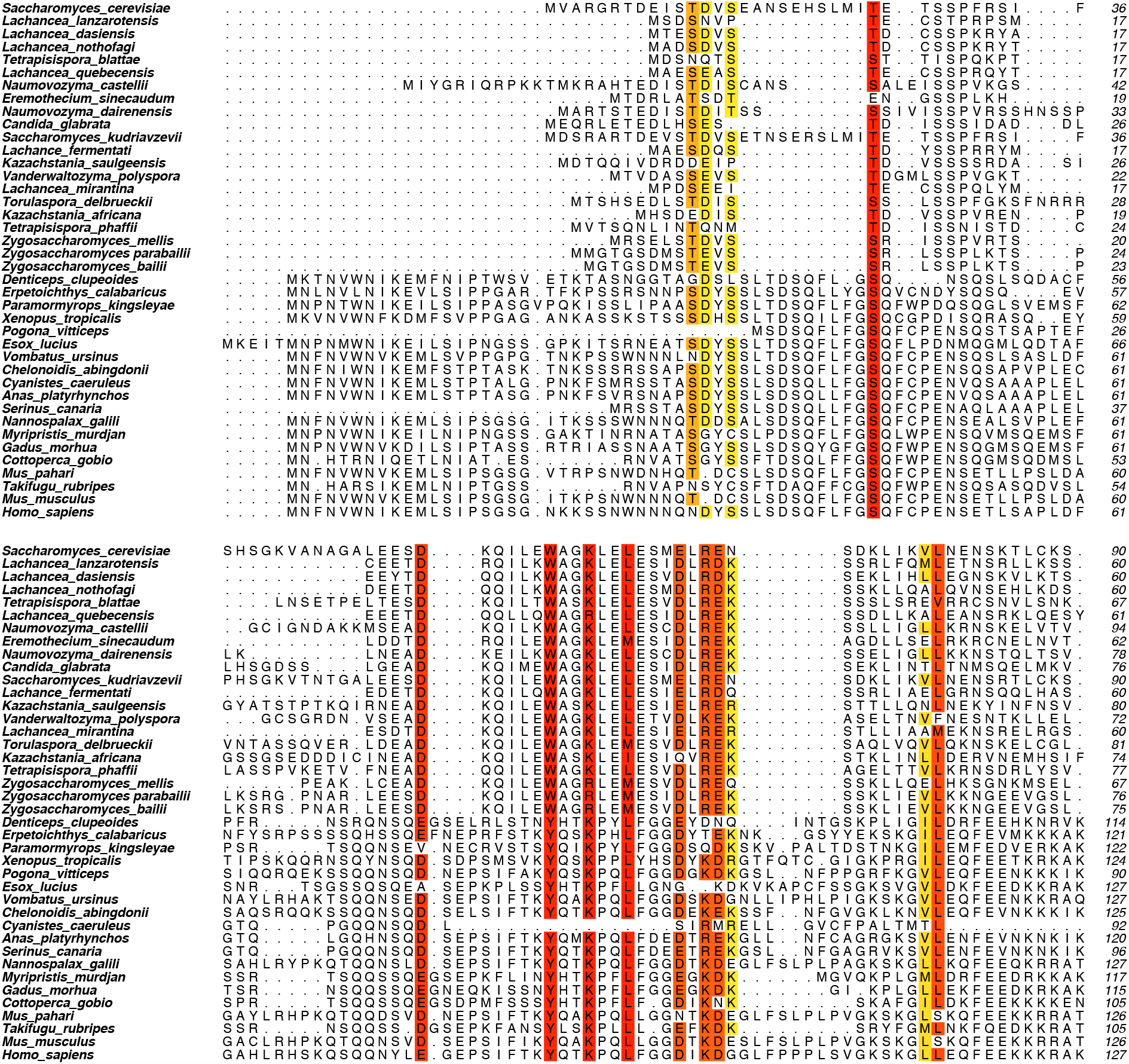
Mer2 sequence alignment An iterative round of BLAST searches and MUSCLE alignments, using Mer2 and IHO1 as starting points were used to generate an alignment of Mer2 orthologs. Residues were coloured from Red to Yellow according to degree of similarity within set similarity groups (DE, FWY, HKR, ILMV, NQ, ST)

## References

1. Keeney, S., Giroux, C. N. & Kleckner, N. Meiosis-specific DNA double-strand breaks are catalyzed by Spo11, a member of a widely conserved protein family. Cell 88, 375–384 (1997).

2. Hunter, N. Meiotic Recombination: The Essence of Heredity. Cold Spring Harb. Perspect. Biol. 7, (2015).

3. Smith, A. V. & Roeder, G. S. The yeast Red1 protein localizes to the cores of meiotic chromosomes. J. Cell Biol. 136, 957–967 (1997).

4. Engebrecht, J. A., Voelkel-Meiman, K. & Roeder, G. S. Meiosis-specific RNA splicing in yeast. Cell 66, 1257–1268 (1991).

5. Engebrecht, J., Hirsch, J. & Roeder, G. S. Meiotic gene conversion and crossing over: their relationship to each other and to chromosome synapsis and segregation. Cell 62, 927–937 (1990).

6. Matos, J. et al. Dbf4-dependent CDC7 kinase links DNA replication to the segregation of homologous chromosomes in meiosis I. Cell 135, 662–678 (2008).

7. Wan, L. et al. Cdc28-Clb5 (CDK-S) and Cdc7-Dbf4 (DDK) collaborate to initiate meiotic recombination in yeast. Genes Dev. 22, 386–397 (2008).

8. Murakami, H. & Keeney, S. Temporospatial coordination of meiotic DNA replication and recombination via DDK recruitment to replisomes. Cell 158, 861–873 (2014).

9. Sommermeyer, V., Béneut, C., Chaplais, E., Serrentino, M. E. & Borde, V. Spp1, a Member of the Set1 Complex, Promotes Meiotic DSB Formation in Promoters by Tethering Histone H3K4 Methylation Sites to Chromosome Axes. Mol. Cell 49, 43 54 (2013).

10. Acquaviva, L. et al. The COMPASS Subunit Spp1 Links Histone Methylation to Initiation of Meiotic Recombination. Science 339, 215 218 (2013).

11. Miller, T. et al. COMPASS: a complex of proteins associated with a trithorax-related SET domain protein. Proc. Natl. Acad. Sci. U. S. A. 98, 12902–12907 (2001).

12. He, C. et al. Structural basis for histone H3K4me3 recognition by the N-terminal domain of the PHD finger protein Spp1. Biochem. J. 476, 1957–1973 (2019).

13. Adam, C. et al. The PHD finger protein Spp1 has distinct functions in the Set1 and the meiotic DSB formation complexes. PLoS Genet. 14, e1007223 (2018).

14. Karányi, Z. et al. Nuclear dynamics of the Set1C subunit Spp1 prepares meiotic recombination sites for break formation. J. Cell Biol. 217, 3398 3415 (2018).

15. Rockmill, B., Engebrecht, J. A., Scherthan, H., Loidl, J. & Roeder, G. S. The yeast MER2 gene is required for chromosome synapsis and the initiation of meiotic recombination. Genetics 141, 49–59 (1995).

16. Kariyazono, R., Oda, A., Yamada, T. & Ohta, K. Conserved HORMA domain-containing protein Hop1 stabilizes interaction between proteins of meiotic DNA break hotspots and chromosome axis. Nucleic Acids Res. (2019)

17. Stanzione, M. et al. Meiotic DNA break formation requires the unsynapsed chromosome axis-binding protein IHO1 (CCDC36) in mice. Nat. Cell Biol. 18, 1208–1220 (2016).

18. n Altmannova, V., Blaha, A., Astrinidis, S., Reichle, H. & Weir, J. R. InteBac - An integrated bacterial and baculovirus expression vector suite. bioRxiv 2020.07.09.194696 (2020)

19. Claeys Bouuaert, C. et al. DNA-driven condensation assembles the meiotic DNA break machinery. Nature (2021)

20. Bouuaert, C. C., Pu, S., Wang, J., Patel, D. J. & Keeney, S. DNA-dependent macromolecular condensation drives self-assembly of the meiotic DNA break machinery. bioRxiv 2020.02.21.960245 (2020).

21. Simon, M. D. et al. The site-specific installation of methyllysine analogs into recombinant histones. Cell 128, 1003–1012 (2007).

22. Davey, C. A., Sargent, D. F., Luger, K., Maeder, A. W. & Richmond, T. J. Solvent mediated interactions in the structure of the nucleosome core particle at 1.9Åresolution. J. Mol. Biol. 319, 1097– 1113 (2002).

23. Young, G. et al. Quantitative mass imaging of single biological macromolecules. Science 360, 423–427 (2018).

24. Fried, M. G. & Bromberg, J. L. Factors that affect the stability of protein-DNA complexes during gel electrophoresis. Electrophoresis 18, 6–11 (1997).

25. Lohr, D. & Van Holde, K. E. Yeast chromatin subunit structure. Science 188, 165–166 (1975).

26. Sollner-Webb, B., Camerini-Otero, R. D. & Felsenfeld, G. Chromatin structure as probed by nucleases and proteases: Evidence for the central role of hitones H3 and H4. Cell 9, 179–193 (1976).

27. Kalashnikova, A. A., Porter-Goff, M. E., Muthurajan, U. M., Luger, K. & Hansen, J. C. The role of the nucleosome acidic patch in modulating higher order chromatin structure. J. R. Soc. Interface 10, 20121022 (2013).

28. Panizza, S. et al. Spo11-accessory proteins link double-strand break sites to the chromosome axis in early meiotic recombination. Cell 146, 372–383 (2011).

29. West, A. M. V., Komives, E. A. & Corbett, K. D. Conformational dynamics of the Hop1 HORMA domain reveal a common mechanism with the spindle checkpoint protein Mad2. Nucleic Acids Res. 46, 279 292 (2017).

30. Raina, V. B. & Vader, G. Homeostatic Control of Meiotic Prophase Checkpoint Function by Pch2 and Hop1. Curr. Biol. 30, 4413–4424.e5 (2020).

31. Yang, C., Hu, B., Portheine, S. M., Chuenban, P. & Schnittger, State changes of the HORMA protein ASY1 are mediated by an interplay between its closure motif and PCH2. Nucleic Acids Res. (2020).

32. Rosenberg, S. C. & Corbett, K. D. The multifaceted roles of the HORMA domain in cellular signaling. J. Cell Biol. 211, 745–755 (2015).

33. Vader, G. Pch2(TRIP13): controlling cell division through regulation of HORMA domains. Chromosoma 124, 333–339 (2015).

34. Subramanian, V. V. et al. Chromosome Synapsis Alleviates Mek1-Dependent Suppression of Meiotic DNA Repair. PLoS Biol. 14, e1002369 (2016).

35. Deshong, A. J., Ye, A. L., Lamelza, P. & Bhalla, N. A quality control mechanism coordinates meiotic prophase events to promote crossover assurance. PLoS Genet. 10, e1004291 (2014).

36. West, A. M. V. et al. A conserved filamentous assembly underlies the structure of the meiotic chromosome axis. Elife 8, e40372 (2019).

37. Kim, Y. et al. The chromosome axis controls meiotic events through a hierarchical assembly of HORMA domain proteins. Dev. Cell 31, 487–502 (2014).

38. Thacker, D., Mohibullah, N., Zhu, X. & Keeney, S. Homologue engagement controls meiotic DNA break number and distribution. Nature 510, 241–246 (2014).

39. Arora, C., Kee, K., Maleki, S. & Keeney, S. Antiviral protein Ski8 is a direct partner of Spo11 in meiotic DNA break formation, independent of its cytoplasmic role in RNA metabolism. Mol. Cell 13, 549–559 (2004).

40. Borde, V. The multiple roles of the Mre11 complex for meiotic recombination. Chromosome Res. 15, 551–563 (2007).

41. Johzuka, K. & Ogawa, H. Interaction of Mre11 and Rad50: two proteins required for DNA repair and meiosis-specific double-strand break formation in Saccharomyces cerevisiae. Genetics 139, 1521–1532 (1995).

42. Cardoso da Silva, R. & Vader, G. Getting there: understanding the chromosomal recruitment of the AAA+ ATPase Pch2/TRIP13 during meiosis. Curr. Genet. (2021)

43. Chen, C., Jomaa, A., Ortega, J. & Alani, E. E. Pch2 is a hexameric ring ATPase that remodels the chromosome axis protein Hop1. Proc. Natl. Acad. Sci. U. S. A. 111, E44–53 (2014).

44. Henderson, K. A., Kee, K., Maleki, S., Santini, P. A. & Keeney, S. Cyclin-dependent kinase directly regulates initiation of meiotic recombination. Cell 125, 1321–1332 (2006).

45. Bhagwat, N. R. et al. SUMO is a pervasive regulator of meiosis. Elife 10, e57720 (2021).

46. Boekhout, M. et al. REC114 Partner ANKRD31 Controls Number, Timing, and Location of Meiotic DNA Breaks. Mol. Cell 74, 1053–1068.e8 (2019).

47. Li, J., Hooker, G. W. & Roeder, G. S. Saccharomyces cerevisiae Mer2, Mei4 and Rec114 form a complex required for meiotic double-strand break formation. Genetics 173, 1969–1981 (2006).

48. Joshi, N., Barot, A., Jamison, C. & Börner, G. V. Pch2 links chromosome axis remodeling at future crossover sites and crossover distribution during yeast meiosis. PLoS Genet. 5, e1000557 (2009).

49. Wojtasz, L. et al. Mouse HORMAD1 and HORMAD2, two conserved meiotic chromosomal proteins, are depleted from synapsed chromosome axes with the help of TRIP13 AAA-ATPase. PLoS Genet. 5, e1000702 (2009).

50. Luger, K., Rechsteiner, T. J. & Richmond, T. J. Preparation of nucleosome core particle from recombinant histones. Methods in Enzymology 3–19 (1999)

51. Weir, J. R. et al. Insights from biochemical reconstitution into the architecture of human kinetochores. Nature 537, 249–253 (2016).

52. Pan, D., Brockmeyer, A., Mueller, F., Musacchio, A. & Bange, T. Simplified Protocol for Cross-linking Mass Spectrometry Using the MS-Cleavable Cross-linker DSBU with Efficient Cross-link Identification. Anal. Chem. 90, 10990–10999 (2018).

53. Kuhl, L.-M. et al. A dCas9-Based System Identifies a Central Role for Ctf19 in Kinetochore-Derived Suppression of Meiotic Recombination. Genetics 216, 395–408 (2020).

54. Vader, G. et al. Protection of repetitive DNA borders from self-induced meiotic instability. Nature 477, 115–119 (2011).

55. Grimm, M., Zimniak, T., Kahraman, A. & Herzog, F. xVis: a web server for the schematic visualization and interpretation of crosslink-derived spatial restraints. Nucleic Acids Research vol. 43 W362–W369 (2015).

